# Stacking models of brain dynamics improves prediction of subject traits in fMRI

**DOI:** 10.1101/2023.11.08.566196

**Authors:** B Griffin, C Ahrends, F Alfaro-Almagro, M Woolrich, S Smith, D Vidaurre

## Abstract

Beyond structural and time-averaged functional connectivity brain measures, the way brain activity dynamically unfolds can add important information when investigating individual cognitive traits. One approach to leveraging this information is to extract features from models of brain network dynamics to predict individual traits. However, there are two potential sources of variation in the models’ estimation which will in turn affect the predictions: first, in certain cases, the estimation variability due to different initialisations or choice of inference method; and second, the variability induced by the choice of the model hyperparameters that determine the complexity of the model. Rather than merely being statistical noise, this variability may be useful in providing complementary information that can be leveraged to improve prediction accuracy. We propose stacking, a prediction-driven approach for model selection, to leverage this variability. Specifically, we combine predictions from multiple models of brain dynamics to generate predictions that are accurate and robust across multiple cognitive traits. We demonstrate the approach using the Hidden Markov Model, a probabilistic generative model of brain network dynamics. We show that stacking can significantly improve the prediction of subject-specific phenotypes, which is crucial for the clinical translation of findings.

## 1 Introduction

A major objective in neuroscience is to develop predictive models that aim to discover associations between brain data and subject traits (e.g., clinical or behavioural measures). Accurate and robust predictions are essential if models are to be used for real-world applications, particularly in clinical settings (Dinsdale et al., 2022). In recent years, for instance, there has been a growing interest in using structural MRI to explore the relationship between ageing, cognitive decline, and certain brain disorders (Kaufmann et al., 2019; Shahab et al., 2019), as well as the illness course of psychoses patients (Mourao-Miranda et al., 2012). However, while structural connectivity can be a powerful predictor, it cannot capture the diverse patterns of neural activity and communication from which cognition emerges. For instance, by analysing resting-state functional MRI (rfMRI), several neurological disorders (e.g., epilepsy, Alzheimer’s) have been associated with disruptions in static (time-averaged) functional connectivity (FC) (Kavitha et al., 2019; Pievani et al., 2011). Furthermore, investigating the dynamics of the brain using fMRI has revealed associations between dynamic (time-varying) FC and complex traits, such as intelligence, beyond anatomy or static FC (Vidaurre et al., 2021).

A common approach to examining brain dynamics is the sliding time window approach (Sakoğlu et al., 2010). Features extracted via the application of this method to fMRI data have been used to predict intelligence (Sen & Parhi, 2021), and the disease progression of Alzheimer’s patients (Abrol et al., 2019). Although there are several variants of this approach (Allen et al., 2014; Leonardi & Van De Ville, 2015; Liégeois et al., 2016), the estimation of FC is inherently noisy and can be influenced by factors such as length and placement of the time windows, affecting the model’s ability to detect temporal variations of interest (Mokhtari et al., 2019; Preti et al., 2017).

An alternative approach to examining brain dynamics is to use generative models of brain network dynamics (that are trained on the entire data set) and then extract features that can predict individual subject traits (e.g., fluid intelligence). One example of this is the use of multivariate autoregressive models, which can capture dynamic FC at a temporal resolution of a few seconds, and have been used to predict task-based phenotypes better than static FC (Liégeois et al., 2019). In our study, we employed the Hidden Markov Model (HMM), a probabilistic model of temporal dynamics of amplitude and FC that characterises the data using a discrete number of amplitude and FC states and transitions between them (Vidaurre et al. 2017). By doing so, we aim to leverage a wider and more diverse range of information.

As showed previously, the HMM can be combined with predictive machine learning methods to predict subject traits from fMRI recordings (Ahrends et al., 2023; Vidaurre et al., 2021). To generate predictions, here we used the Fisher kernel method, a mathematically principled way of combining generative models like the HMM with predictive methods through the use of a kernel function (Schölkopf & Smola, 2001; Shawe-Taylor et al., 2004). This approach enabled us to preserve the structure of the generative model while leveraging the predictive power of kernel methods (Ahrends et al., 2023; Jaakkola et al., 2000; Jaakkola & Haussler, 1998).

However, the practical implementation of subject trait predictions based on functional data relies on the accuracy and robustness of the predictions. Regardless of the chosen generative model, there can be two potential sources of variation in these predictions. First, there can be run-to-run variability (e.g., due to different initialisations of the estimation process for a given dataset) for approaches relying on non-convex optimisation procedures (Alonso & Vidaurre, 2023); the HMM inference is an example of this, which can impact subsequent predictions made using features derived from the fitted HMM model. Second, the choice of generative model hyperparameters, such as the model order (e.g., number of HMM states) or the strength of regularisation, affects the result of the estimation, so modifying these hyperparameters can add an additional factor of variability. Variability is generally viewed as a disadvantage because it poses a problem for interpretability. When it comes to predicting subject traits using the HMM, moreover, it can be difficult to determine the optimal HMM run or hyperparameter choice, potentially leading to inconsistent prediction accuracies.

In our study, however, we aimed to identify variability that offers distinct and complementary information (across different analyses of the same data) rather than being driven by estimation noise. Then, we employed a stacked generalisation framework (Breiman, 1996; Wolpert, 1992) that leverages this variability to improve the accuracy and robustness in predicting subject traits. Previous studies have used stacking to integrate multimodal neuroimaging data for age prediction (Engemann et al., 2020) and mild cognitive impairment classification (Vaghari et al., 2022). In this study, we used stacking to substitute model selection with model integration—in other words, our approach bypassed the complex decision of selecting the best HMM hyperparameter choices, albeit at the expense of increased computation. Specifically, our method aimed to integrate complementary descriptions of brain dynamics, such as capturing different temporal scales, as they are captured by different configurations of the HMM (i.e., across a wide range of hyperparameters).

Focusing on resting-state fMRI (rfMRI) due to its wide availability, we demonstrate the efficacy of our stacking framework on data from two large-scale neuroimaging datasets: UK Biobank (UKB) (Sudlow et al., 2015) and the Human Connectome Project (HCP) (Van Essen et al., 2013). For both datasets, we explored the relationships between rfMRI data and a range of cognitive traits. By doing so, we found that stacking predictions generated from different configurations of the HMM generally outperformed simpler approaches in terms of robustness and accuracy.

## 2 Materials & Methods

To predict a set of cognitive traits from models of rfMRI FC dynamics, our approach involved multiple steps. In brief, we first independently ran the HMM on the rfMRI data multiple times, generating a prediction from each HMM (referred to as base-level predictions). Subsequently, we used these base-level predictions as input features for a meta-model to generate a stacked prediction.

The process used to produce a base-level prediction is outlined in **Figure 1**. For each base-level prediction, an HMM was run on a temporally concatenated fMRI timeseries of all subjects to obtain a group-level HMM (Vidaurre et al., 2017, 2021) (**Figure 1a**). We then used the Fisher kernel method, a mathematically principled approach to predicting target variables from an HMM (Ahrends et al., 2023; Jaakkola & Haussler, 1998) (**Figure 1b**). We compared each subject’s timeseries to the group-level HMM in the HMM’s parameter space, assuming that similar subjects would induce similar parameter gradients in this space. Using these comparisons, we created a subjects-by-subjects similarity matrix—the Fisher kernel matrix. Finally, subject-specific trait predictions were generated using these similarity matrices via a kernel ridge regression (KRR) model (**Figure 1c**).

**Figure 1.**
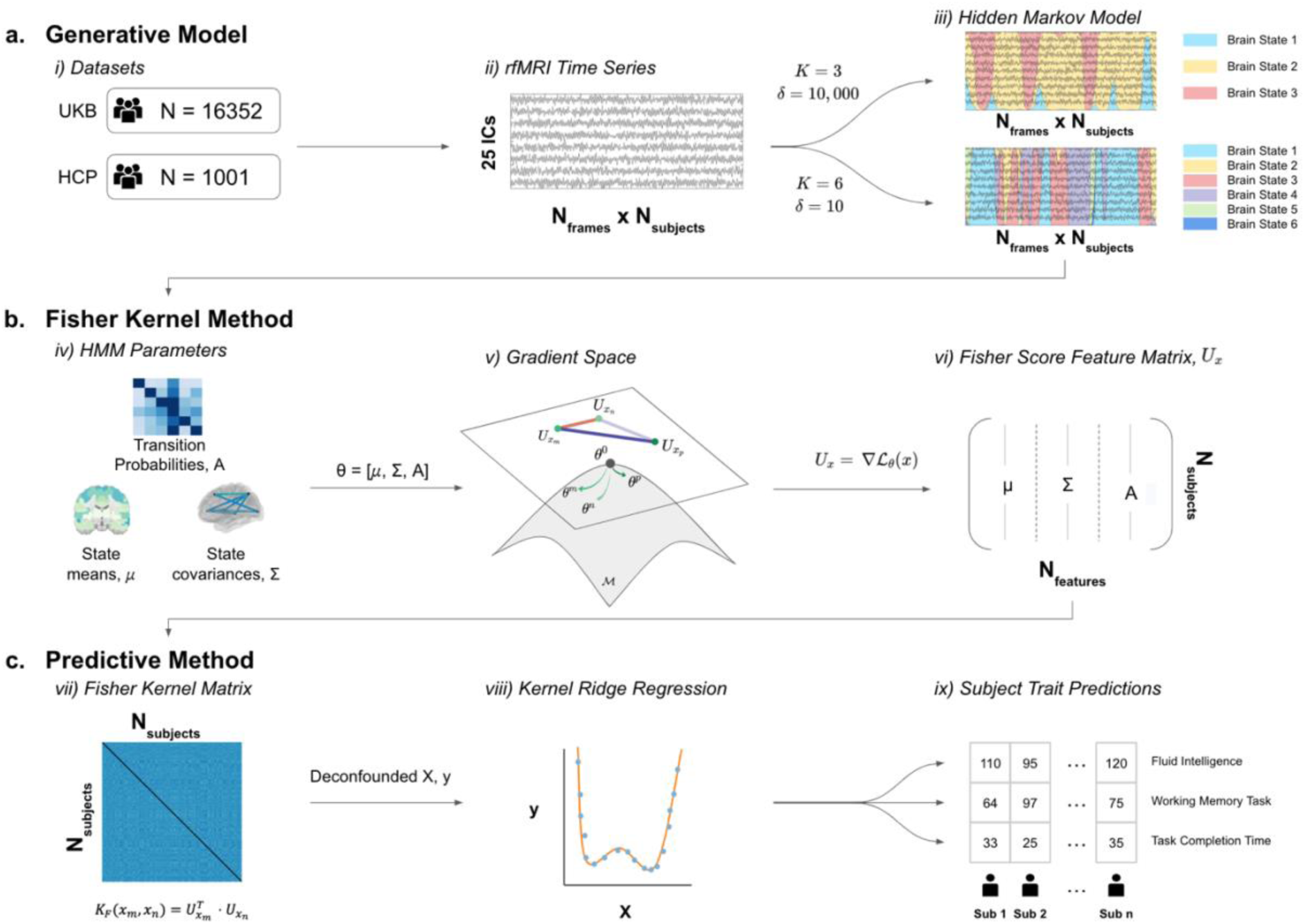
Procedure for predicting subject traits from resting-state fMRI (rfMRI) timeseries. **(a) Generative Model.** (i), (ii) rfMRI in groupICA parcellations with 25 ICs are concatenated across all subjects from UKB and HCP, respectively. (iii) The hidden Markov model is trained on the timeseries, where different HMM hyperparameters, such as number of states (*K*) or the prior probability of remaining in the same state (*δ*), lead to different HMM descriptions. **(b) Fisher Kernel Method.** (iv) The HMM consists of state-time courses and the parameters defining the model (*θ*), including the brain state means (*μ*) and covariances (*Σ*), and the transition probability matrix (*A*). (v) Fisher scores are generated for each subject’s parameter set by taking the derivative of the log-likelihood with respect to each parameter to determine those which are instrumental in forming the subject-specific state-time courses. (vi) The Fisher scores are vectorised for each subject and combined to form the Fisher score feature matrix. **(c) Predictive Method.** (vii) We then construct the practical Fisher kernel from all subjects’ Fisher scores to form a subject-by-subject similarity matrix—the Fisher kernel matrix. (viii) The Fisher kernel is used as a predictor in a (nested) cross-validated, deconfounded kernel ridge regression model to predict subject traits, where the optimal regularisation parameters are found via 10-fold cross-validation. (ix) This last step is carried out independently to predict multiple cognitive traits.

Specifically, we generated 100 base-level predictions in this way, independently estimating an HMM for each base-level prediction from the same fMRI timeseries; 50 times with fixed hyperparameters (to investigate the run-to-run variability of the HMM), and 50 times with varying hyperparameters (to investigate variability induced by varying the hyperparameters). We then generated two stacked predictions by separately combining the two sets of 50 predictions. We now present all the steps that integrate this procedure in more detail and introduce the two datasets to which we applied the framework, UKB and HCP.

### 2.1 HCP neuroimaging and non-imaging data

We used publicly available rfMRI data from 1,001 subjects from the Human Connectome Project (HCP) dataset (Van Essen et al., 2013). The full acquisition details and preprocessing pipeline for the HCP dataset are described in Van Essen et al. (2012). In brief, 3T whole-brain fMRI data were acquired with a spatial resolution of 2×2×2 mm and a temporal resolution of 0.72s. The preprocessing pipeline is described in (Smith et al., 2013), but the primary steps included motion correction, high-pass temporal filtering to remove the linear trends of the data, and artefact removal using ICA+FIX (Griffanti et al., 2014). A parcellation of 25 independent components (ICs) was obtained by performing group spatial ICA using MELODIC (Beckmann & Smith, 2004). We chose the 25 IC parcellation as it covers the major functional networks. Additionally, coarser parcellations are more reliable for estimating FC dynamics due to the smaller number of free parameters per state (Ahrends et al., 2022). For each participant, this resulted in 25 timeseries, composed of 4,800 time points across four scanning sessions (with 1,200 time points in each session) of approximately 15 minutes each. For performance evaluation, we selected all traits related to fluid intelligence, as well as the traits given by the unadjusted cognitive test scores. This resulted in 15 cognitive traits. The list of these 15 traits can be found in **Supplementary Table SI-1**.

As in Vidaurre et al. (2017), we controlled for sex and motion. Additionally, we accounted for familial relationships when assigning subjects to cross-validation folds, to make sure related subjects were never split across folds.

### 2.2 UKB neuroimaging and non-imaging data

Our aim was to predict a set of cognitive traits (e.g., task performance) for 16,352 subjects from the UK Biobank (UKB) dataset (Sudlow et al., 2015) using models of their rfMRI dynamics. The full acquisition details and preprocessing pipeline for the UKB dataset are described in Smith et al. (2022). In brief, 3T fMRI data were acquired for each participant, consisting of 490 time points per session at a TR of 0.735s and with 2.4mm spatial resolution. Preprocessing was performed using the standard UKB pipeline, including brain extraction, motion correction, structured artefact removal using ICA+FIX (Griffanti et al., 2014), high-pass temporal filtering and registration to MNI152 space (Alfaro-Almagro et al., 2018). We used surface-based node timeseries generated using 25-dimensional group-ICA surface maps from the Human Connectome Project (HCP) S1200 dataset (Van Essen et al., 2013). This resulted in 25 timeseries, composed of 490 time points across a 6-minute scanning session for each patient, and allowed for a more direct comparison between UKB and HCP.

To match the number of traits investigated in HCP, we first excluded cognitive traits with missing recorded values for over half of UKB subjects, before performing preliminary analyses to generate predictions for the remaining 450 cognitive traits in UKB (of the 1331 cognitive traits in total). Subsequently, we narrowed down our focus to the top-performing 15 traits in terms of prediction accuracy. The rationale behind selecting the traits which could be predicted most accurately was to demonstrate the potential of stacking compared to alternative approaches. Since stacking involves combining predictions from base estimators, these underlying estimators must exhibit a certain level of accuracy for stacking to be any meaningful. Given that the primary focus of our investigation was to compare predictions from dynamic FC, this preliminary analysis was performed using static FC to reduce bias towards a particular dynamic FC approach. The prediction accuracies for all 450 cognitive traits are shown in **Supplementary Figure SI-1** and the list of the selected 15 traits can be found in **Supplementary Table SI-2**.

Also, we employed a reduced set of confounds extracted from a comprehensive set of 602 UKB imaging confounds provided by Alfaro-Almagro et al. (2021). To generate our reduced set of confounds, we first selected a subset of conventional confounds including sex, scanning site, head size, and head motion (which use FSL’s FEAT (Woolrich et al., 2001) and EDDY (Andersson et al., 2016, 2017)). Then, we reduced the remaining confounds by using the singular value decomposition and selecting the top principal components which accounted for 85% of the variance. This process resulted in a set of 30 variables that we used for deconfounding by regressing them out from both the predictor and target variables using linear regression. To prevent information leakage from the test data to the training data, we applied deconfounding within cross-validation folds.

### 2.3 The hidden Markov model

The hidden Markov model is a generative probabilistic model that can be applied to fMRI timeseries to find recurrent patterns of activity and FC (Vidaurre et al., 2017). The model assumes that the data can be described using a discrete number of probabilistic models, or latent states. While it is possible to parametrise the HMM to focus primarily on FC, here we chose to model both FC and signed amplitude for the majority of our analysis, but also considered models with only FC (see **Supplementary Figure SI-2**).

An adequate choice for fMRI is to model these brain states as Gaussian distributions, where each state *k* (of a total *K* states) is governed by two sets of parameters - a mean vector *μ*_*k*_, which can be interpreted as the mean activation for each of *M* brain regions from a specified parcellation (here *M* group-ICA components), and a covariance matrix Σ_k_, which can be interpreted as FC between the *M* brain regions. The model inference also estimates the initial probabilities that the timeseries start in each of the *K* states, given by *π*, as well as the probabilities of transitioning between states, given by a transition probability matrix *A*.

The HMM used to represent the fMRI data is described by this set of parameters, *θ* (see **Figure 1a**), where *μ* and Σ represents the mean vectors and covariance matrices across all states:

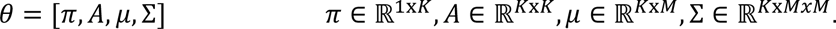

The optimal parameters are estimated from the data at the group level (i.e., the state probability distributions are the same for all subjects)^1^. Following this, subject-specific HMM parameters can be determined using dual estimation (Vidaurre et al., 2021), as well as subject-specific state-time courses (i.e., the probabilities of each state being active at each time point).

In the HMM generative process, we assume that each time step *t* of the observed fMRI timeseries *X*_*t*_ involves sampling from a Gaussian distribution with mean *μ*_*k*_ and covariance Σ_*k*_ when state *k* is active:

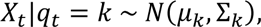

where *q*_*t*_ is the active state at time *t*. This currently active state, *q*_*t*_, depends on the previous state *q*_*t*−1_ and is determined by the transition probabilities *A*. Consequently, the sequence of states is generated by sampling from a categorical distribution with parameters:

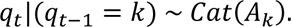

where, *A*_*k*_ represents the *k*-th row of the transition probability matrix.

When applying the HMM, there are two potential sources of variability. The first we refer to as run-to-run estimation variability, which can arise both from the random initialisation of the HMM estimation process for a given dataset, as well as from the stochastic inference method we apply. Random initialisation refers to the random assignment of parameter values, such as the state parameters or transition probabilities, at the beginning of the training process. Furthermore, we use an efficient stochastic approach for the HMM estimation, which applies the principles of stochastic optimisation to variational inference (Vidaurre et al., 2018). This approach is computationally cheaper and suitable for large neuroimaging datasets such as UKB and HCP. However, it introduces variability in the estimation of the state-time courses, which are determined by the Baum-Welch algorithm (Baum et al., 1970). At each iteration, the estimation of the state-time course is updated based on a random subset of subjects, which is a noisy but computationally efficient approach. Since the HMM optimisation can converge to local minima, different initialisations and batches of subjects for the estimation can impact the accuracy of subsequent predictions.

The second source of variability comes from the selection of the hyperparameters of the HMM, which are set by the user. We refer to this as hyperparameter selection variability. By varying the hyperparameters, we can discover distinct patterns of FC, for example across different time scales (Ahrends et al., 2022). In our study, we focused on varying two HMM hyperparameters: the number of states (*K*), and the prior probability of remaining in the same state (δ), which is parametrised by the prior Dirichlet distribution concentration parameter of the corresponding prior distribution^2^ (Masaracchia et al., 2023), which effectively influences the time scale of the estimate. This parameter determines the size of the diagonal elements in the prior distribution of the hidden states (i.e., the transition probability matrix).

In total, we fit the HMM to the same rfMRI timeseries 100 times; 50 times with fixed hyperparameters^3^ (i.e., with random initialisations and random batches of subjects for the stochastic inference;), and 50 times with varying hyperparameters^4^ (i.e., with additional variability coming from the selection of hyperparameters;). When fixing the hyperparameters, we chose *δ* = 10, the default setting in the HMM-MAR toolbox, and *K* = 6, commonly used values in recent literature (Alonso & Vidaurre, 2023; Quinn et al., 2018; Vidaurre et al., 2018). The choice of states represents a compromise between having a higher number of states, which can increase the chances of certain states being present in only a subset of subjects (Ahrends et al., 2023), and a lower number of states, which can increase the chance of assigning entire sessions to a single state (i.e., model stasis) (Ahrends et al., 2022). Nevertheless, the specific choice of hyperparameters is less critical to our study, as our focus when fixing the hyperparameter is to investigate run-to-run variability which could be done with an alternative configuration.

When we varied the hyperparameters, our objective was to investigate both run-to-run variability and hyperparameter selection variability, irrespective of the specific hyperparameter settings employed. Therefore, we used a wide range of hyperparameters and repeated each hyperparameter combination twice with different initialisations and different subject batches for the stochastic inference. As a result, we anticipated a greater degree of variability when adjusting the hyperparameters of the HMM, since we expect to encounter both types of variability.

#### 2.3.1 Static FC

To compare the performance of our dynamic FC predictions with simpler methods that do not consider brain dynamics, we also used time-averaged FC (also referred to as static FC) for prediction. To be comparable to the dynamic FC predictions, we used a one-state HMM as a static FC estimator, which essentially yields a single covariance matrix per subject (modelled as an inverse Wishart distribution). Subsequently, we used the Fisher kernel method and kernel ridge regression in an analogous manner to the dynamic FC predictions.

Alternatively, it is also possible to use partial correlations, which is defined as the correlation between the time series of two brain regions after regressing out the time series of all the other regions (i.e., the inverse covariance). We repeated our static FC analysis using this alternative approach, the results of which can be found in **Supplementary Figure SI-3.**

### 2.4 The Fisher kernel

To predict subject traits from the HMM, we used the Fisher kernel method (Ahrends et al., 2023; Jaakkola et al., 2000; Jaakkola & Haussler, 1998), which is a mathematically principled approach to predict from generative probabilistic models. To do this, we first estimate subject-specific HMM parameter sets (i.e., initial state probabilities, transition probability matrices, and brain state means and covariance) using dual estimation (Vidaurre et al., 2021). The Fisher kernel is constructed from these subject-specific HMM parameter sets and can be used with a kernel-based classification or prediction method, here KRR, while respecting the structure of the HMM. Specifically, the Fisher kernel is constructed upon the Riemannian manifold formed by the parameters that define the HMM (Ahrends et al., 2023; Jaakkola et al., 2000; Jaakkola & Haussler, 1998). This method therefore provides an approach, given a certain choice of hyperparameters, to predict subject traits from time-varying patterns of activity and FC.

The intuition behind the Fisher kernel is in the comparison of the generative processes for the subjects’ timeseries. Specifically, we look at the influence of a particular HMM parameter in the generation for the timeseries of each subject and compare these for every pair of subjects. This is represented by the Fisher scores, which are gradients in the parameter space. Fisher scores are then compared between subjects, such that similar subjects have similar scores for fixed HMM parameters.

The Fisher score, *U*_*x*_, is given by the gradient of the log-likelihood with respect to each parameter (**Figure 1b**):

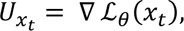

where *x*_*t*_ is the fMRI timeseries of a given subject, and ℒ_*θ*_(*x*_*t*_) = *P*(*x*_*t*_|*θ*) is the log-likelihood of timeseries *x*_*t*_ given the HMM parameters *θ*.

The Riemannian manifold upon which the parameters of the HMM lie is completed by choosing a local Riemannian metric, with the Fisher information matrix being a natural choice for probability distributions (Amari & Nagaoka, 2000). The Fisher information matrix, ℱ, is defined as:

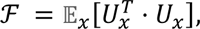

where the expectation is taken with respect to *x* under the distribution *p*(*x*|*θ*).

The invariant Fisher kernel, *K*_*I*_, is then defined as the inner product of Fisher scores, *U*_*x*_, scaled by the Fisher information matrix:

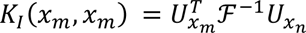

for subjects *m* and *n*, where the Fisher information matrix accounts for the different scales of the various model parameters.

In practice, the Fisher information matrix is often disregarded because its empirical approximation is simply the covariance matrix of the Fisher scores, meaning that using it would be equivalent to whitening the scores. This makes its impact negligible as sample size increases while being computationally expensive (Jaakkola & Haussler, 1998; Shawe-Taylor et al., 2004). As a result, the invariant Fisher kernel is frequently reduced to the practical Fisher kernel, *K*_*F*_, for which the linear version is defined as:

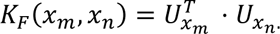

The dot product is computed for all subject pairs to obtain a similarity matrix, or kernel matrix, which can be used with any suitable kernel prediction method or classifier. In our study, we adopted a linear kernel in combination with KRR.

### 2.5 Kernel ridge regression (KRR)

After developing a kernel matrix, kernel methods can be used to determine the relationship between a subject’s time-varying activity and FC patterns and their cognitive traits.

In this study we used KRR due to its prevalence in neuroimaging literature, efficiency, and ability to achieve comparable performance to more complex approaches such as deep learning (He et al., 2020; Mihalik et al., 2019).

KRR is the kernelised version of ridge regression (Saunders et al., 1998). Given a subject trait to predict, *y*, regression coefficients, *α*, are determined by solving the optimisation problem:

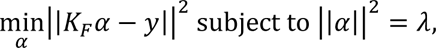

where *λ* is the L2-regularisation chosen by cross-validation and *K*_*F*_ is the practical Fisher kernel. The optimal regression coefficients, *α*^∗^, can then be estimated as:

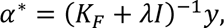

where *I* is the identity matrix.

We used KRR to generate base-level predictions corresponding to independent runs of the HMM on the same data, for 15 cognitive traits from the UKB and HCP datasets.

### 2.6 Stacking

Stacking, or stacked generalisations, is a method which can be employed to potentially improve the predictions generated from methods such as KRR. Specifically, stacking is a technique that combines the predictions from multiple base models by training a meta-model that uses the base model predictions as input features.

Given multiple base models, the objective is to determine optimal model coefficients, or stacking weights, for combining the base-level predictions to generate the best stacked predictions for a given subject trait. To achieve this, a common approach is to use the base-level predictions as input features in a constrained linear regression, where the model coefficients are forced to be non-negative and sum to 1 (Breiman, 1996):

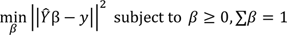

Where 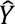 is the feature matrix (formed of base-level predictions), *y* represents the corresponding subject traits, and *β* are the stacking weights to be determined.

The base-level predictions are likely to be highly correlated, particularly when generated from HMMs with identical hyperparameters. Therefore, if unconstrained least squares is used to determine the stacking weights, there is no guarantee that the resulting stacked prediction would stay within the range of the base-level predictions and generalisation may be poor (Breiman, 1996). However, by imposing non-negativity and sum-to-1 constraints, an “interpolating” prediction is developed (i.e., the stacked prediction remains within the range of the base-level predictions).

The primary contribution of this paper is the implementation of a stacked generalisation scheme (Wolpert, 1992) to combine predictions obtained from multiple HMM runs using the Fisher kernel method. After developing multiple base-level predictions, we used a two-layered nested cross-validation scheme to generate out-of-sample stacked predictions (i.e., predictions for unseen subjects).

A summary of the stacking and nested cross-validation framework is depicted in **Figure 2**. Initially, the subjects were divided into 10 folds. Each fold was sequentially chosen as the outer loop test set, while the remaining 9 folds formed the outer loop training set. This outer loop served as the first layer of the nested cross-validation and was used to evaluate the performance of the stacked model. The subjects in the outer loop training set (i.e., 90% of the full sample) were further divided into 10 folds, which served as the 9 training folds and 1 test fold for the inner loops of the cross-validation. This constituted the second layer of the nested cross-validation, involving two inner loops using the same cross-validation folds. These inner loops served two purposes: optimising the regularisation parameter and determining stacking weights.

**Figure 2.**
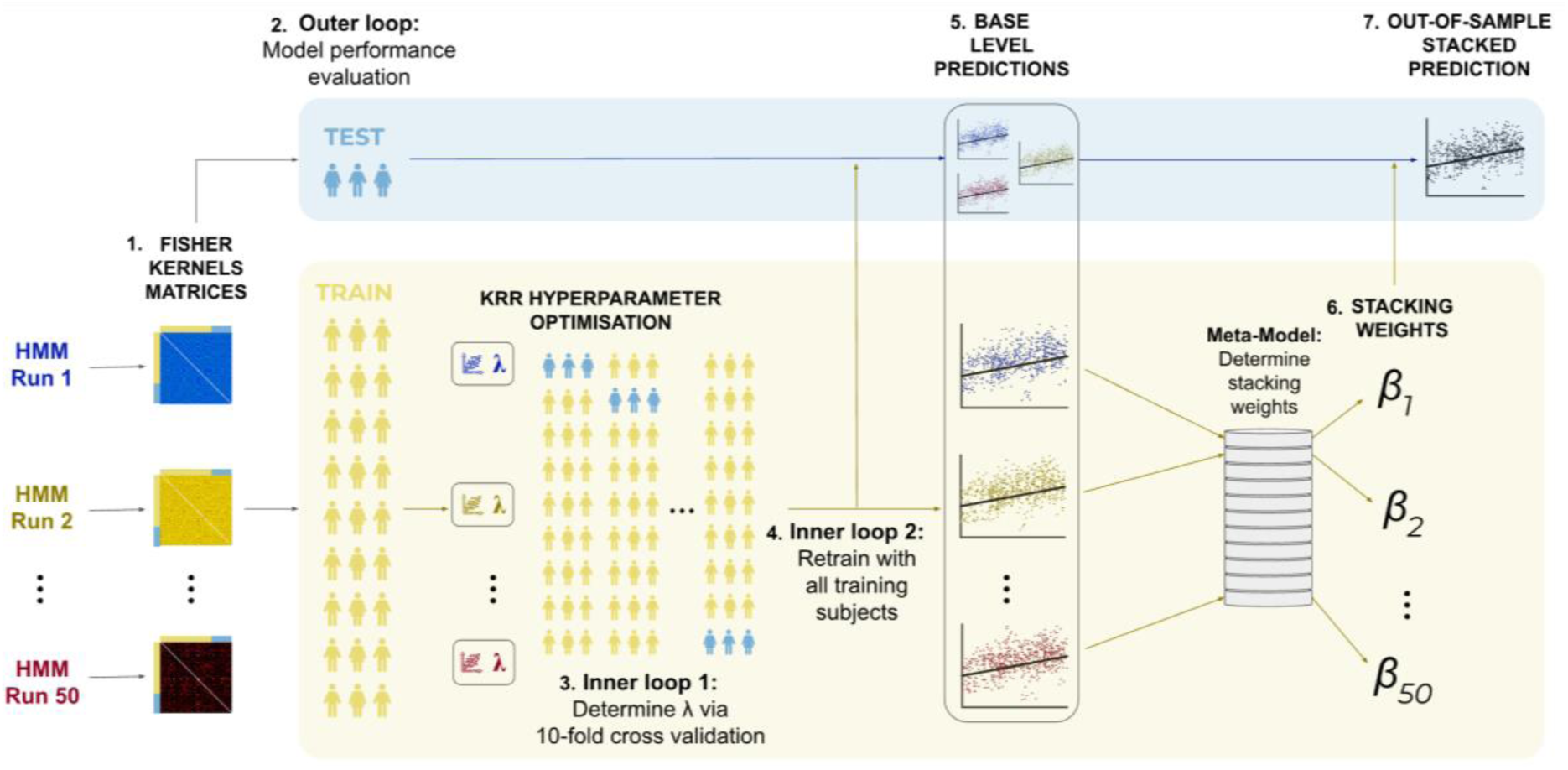
Cross-validation framework for stacking 50 base-level predictions developed from Fisher kernels generated from the features of 50 distinct hidden Markov models. **(1)** 50 group-level HMMs were run on the concatenated rfMRI timeseries across subjects, from which subjects-by-subjects similarity matrices were developed by applying the Fisher kernel method to each HMM. **(2) Outer loop:** the subjects were split into 10 cross-validation folds, where each fold was selected in turn as an outer loop test set (used for stacked model performance evaluation), and the remaining 9 folds are used as an outer loop training set. The training subjects were further divided into 10 cross-validation folds, which are used for two purposes. **(3) Inner loop 1:** this loop was used to optimise the L2-regularisation parameter, *λ*. **(4) Inner loop 2:** using the optimised *λ* from inner loop 1, base-level predictions were developed for all inner loop subjects (or equivalently the outer loop training subjects). **(5)** Separately, base-level predictions are generated for the outer loop test subjects using the optimised hyperparameter from **inner loop 1**. **(6)** The 50 base-level estimators from **inner loop 2** were then stacked, with the stacking weights constrained to be non-negative and sum to 1. **(7)** Finally, the stacking weights are combined with the base-level predictions for the outer loop test subjects to produce out-of-sample stacked predictions for a selected cognitive trait. This is repeated for all 10 outer loop cross-validation folds.

The first inner cross-validation loop (see *inner loop 1* in **Figure 2**) was used to tune the L2- regularisation parameters of the KRR models. This process was carried out separately for each Fisher kernel matrix obtained from the respective HMM. Following that, the second inner cross-validation loop, (see *inner loop 2* in **Figure 2**) was used to determine stacking weights. In this step, the optimised KRR regularisation parameters (from inner loop 1) were used to generated 50 base-level predictions for all inner loop subjects (or, equivalently, the outer loop training subjects; see *inner loop 2* in **Figure 2**). These base-level predictions were subsequently used as input features in a meta-model to generate stacking weights, where the stacking weights were forced to be non-negative and sum to 1.

Finally, we developed stacked predictions for the outer loop test set. To do this, we used the optimised KRR regularisation parameters generated in inner loop 1 to generate base-level predictions for the outer loop test set. We then combined these outer loop test set predictions with the stacking weights generated from inner loop 2 (in other words the outer loop training set) to produce out-of-sample test set stacked predictions, used to evaluate model performance. We completed this analysis for 50 HMMs with fixed hyperparameter (to test run-to-run variability of the HMM), and 50 HMMs with varying hyperparameters (to test hyperparameter selection variability). This resulted in two sets of 50 base-level predictions and two corresponding stacked predictions. Through this approach, we sought to create a more robust stacked prediction with superior accuracy.

By varying the hyperparameters of the HMM, our objective was to explore different configurations that could reveal unique brain state distributions, accurately capturing information across subjects. However, adjusting these hyperparameters can sometimes lead to inadequate characterisation of brain data. For instance, the choice of the number of states can impact model stasis (Ahrends et al., 2022). To assess the diversity of the predictions, we examined the correlation between predictions generated by distinct HMMs to indicate if they potentially contained differential information. Additionally, we examined the accuracy of these predictions to discern whether diversity was driven by inaccurate predictions, or whether they were distinct and complementary. By combining diverse yet accurate predictions (e.g., multiple predictions which are accurate but each one explains one aspect of the data better than the others), we aimed to develop accurate subject-specific trait predictions. This approach also offers the advantage of circumventing the challenge of model selection by combining the information from multiple representations of the data.

### 2.7 Performance Evaluation

We evaluated predictions for 16,352 subjects from UKB, and 1,001 subjects from HCP, or as many subjects as data were available for a given trait. To investigate the effectiveness of stacking, we compared the performance of the stacked prediction with the base-level predictions that were combined to generate it. We also compared our stacked prediction with the common and most simplistic form of combination, which involves taking the average of all base-level predictions so that each one contributes equally to the final prediction (Lincoln & Skrzypekt, 1989; Perrone & Cooper, 1992; Tumer & Ghosht, 1996). In other words, given *k* base-level predictions, 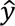_*k*_, we combine them as follows:

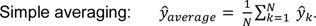

Our models were assessed based on accuracy and robustness. Accuracy was measured using the coefficient of determination, R^2^. Given an observed subject trait, *y*, and our prediction, 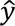, for *N* subjects, the coefficient of determination is given by

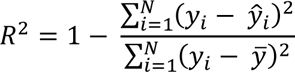

where 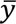 is the mean of the observed subject trait over all subjects.

For a model to be robust, the model’s accuracy should be consistent across variations of the training set. Robustness is extremely important for real-world applications, but often a short-coming in neuroimaging-based prediction studies (Varoquaux et al., 2017).

We estimated prediction accuracy using 10-fold cross-validation. For each trait, we repeated the cross-validation process 10 times using different randomised folds, following the approach of Džeroski & Ženko (2004). This allowed us to assess the consistency and variance of our models’ accuracy. To assess accuracy, we calculated the mean R^2^ scores across the 10 cross-validation iterations. To assess robustness, we examined the variance of the R^2^ scores across the 10 iterations.

To assess the statistical significance of the differences in accuracy between our different approaches, we used Welch t-tests. Specifically, we compared the mean prediction accuracies across 15 subject traits and 10 cross-validation iterations for each dataset separately. We used Welch’s t-test since the variance between the groups compared was different between the base-level predictions from HMMs with fixed versus varying hyperparameters for certain traits (see **Supplementary Table SI-4** for details). To account for multiple comparisons (i.e., the 30 different tests composed of the 15 comparisons between different stacking approaches for the 2 datasets), we applied Benjamini-Hochberg’s False Discovery Rate (FDR) procedure (Benjamini & Hochberg, 1995) to correct the resulting p-values.

To investigate the robustness of stacking, we compared the variance of the distribution of accuracies across cross-validation iterations between base-level predictions and stacked predictions. Since the distributions did not follow a normal distribution for certain traits (see **Supplementary Table SI-5** for details), we used Levene’s test to compare the distributions.

Similar to our previous analysis, p-values were corrected across using Benjamini-Hochberg’s FDR procedure (across the 60 different tests comprised of 15 from UKB and 15 from HCP tests for the two sets of Levene’s tests).

## 3 Results

In this section, we investigate the effectiveness of stacking predictions using HMMs with fixed and HMMs with varying hyperparameters. We find that stacking predictions from HMMs with fixed hyperparameters generally exhibits moderate effectiveness. On the other hand, stacking predictions from HMMs with varying hyperparameters substantially improves accuracies. Although stacking performs similarly to simple averaging in many situations, we find that there are certain scenarios where stacking yields much higher accuracies. Furthermore, we show that stacking produces robust predictions, particularly when combining predictions from HMMs with varying hyperparameters. We then explore the factors contributing to the effectiveness of stacking in certain scenarios. Finally, we showcase the flexibility and power of stacking by combining predictions from static FC with those from dynamic FC.

### 3.1 Stacking predictions generated from HMMs with fixed hyperparameters improves prediction accuracy only moderately

Our analysis revealed that combining predictions generated from 50 HMMs with fixed hyperparameters (i.e., those with run-to-run variability caused by different initialisations and the stochasticity of the HMM inference) resulted in a higher average prediction accuracy than base-level predictions for UKB but was only effective for certain subject traits in HCP. **Figure 3** provides a comparison of the distribution of explained variance (coefficients of determination; R^2^) between predictions generated from HMMs with fixed hyperparameters (depicted in blue) and observed subject traits for three approaches: base-level predictions (shown by the boxplots), averaging base-level predictions (shown by the crosses), and stacking them using constrained least squares (shown by the triangles).

**Figure 3.**
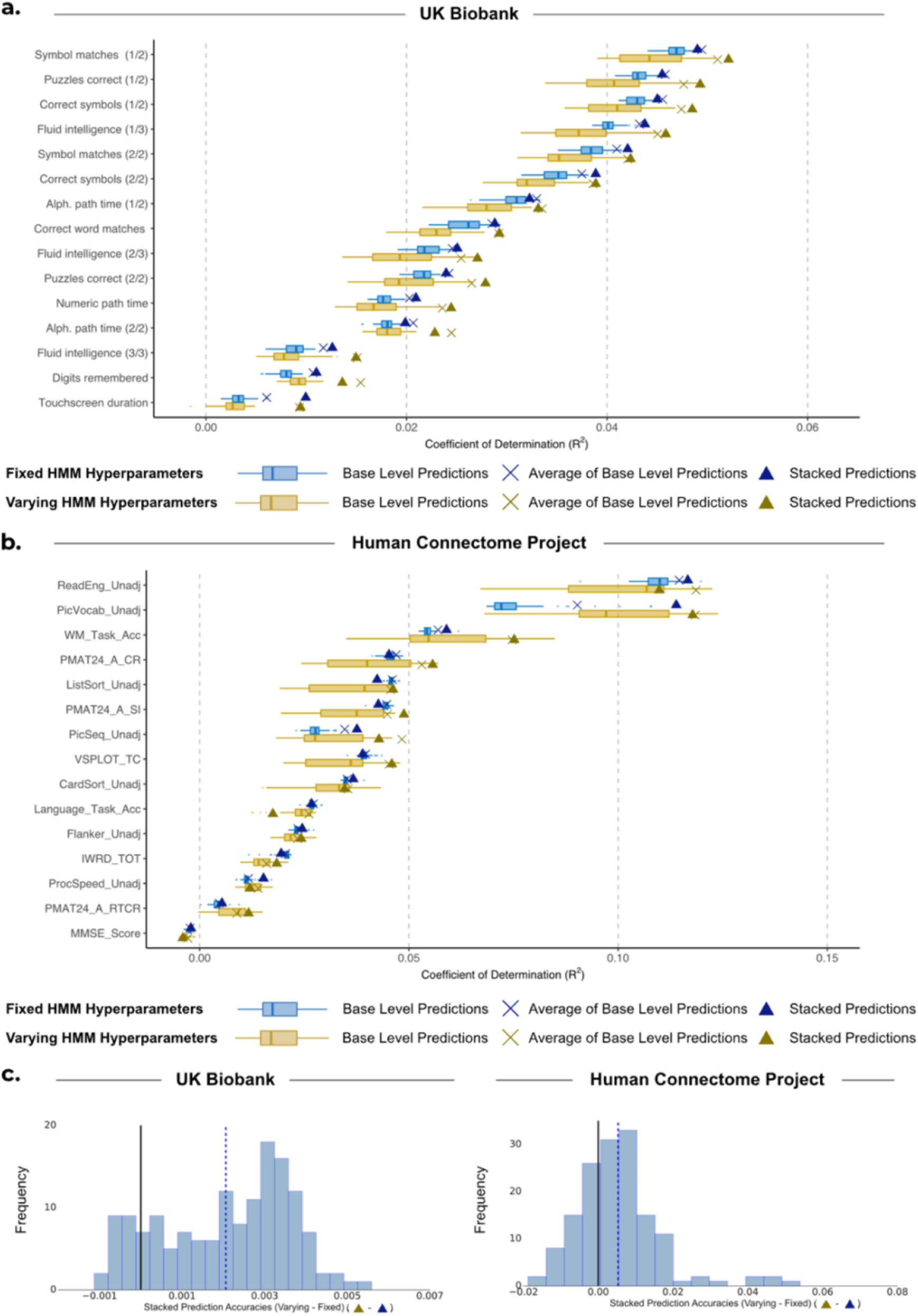
Comparison of performance for base-level predictions and stacking predictions from HMMs with varying hyperparameters against HMMs with fixed hyperparameters. **(a, b)** Performance of stacking across subject traits for UKB and HCP respectively. Boxplots show the R^2^ scores between observed subject traits and base-level predictions generated from 50 HMMs. These are compared to the R^2^ scores when we combine the base-level predictions by taking the average of them (×) and by stacking (Δ). Blue represents the results of using HMMs with fixed hyperparameters. Yellow represents the results of using HMMs with varying model hyperparameters. **(c)** Distribution of the difference between stacking prediction using varying hyperparameters (Δ) and fixed hyperparameters (Δ) across 10 cross-validation

The results for the UKB are depicted in **Figure 3a**. Across 15 traits and 10 cross-validation repetitions, stacking exhibited a significantly higher average prediction accuracy compared to base-level predictions (mean R^2^ _STACKED_: 0.0299 vs. mean R^2^ _BASE_: 0.0268; *p_BH_*=0.0091). Additionally, for 10 out of the 15 traits investigated, the stacked prediction outperformed every base-level prediction. Despite this, stacking performed equally to simply averaging across the base-level predictions (mean R^2^ _STACKED_: 0.0299 vs. mean R^2^ _AVERAGE_: 0.0295; *p_BH_*=0.499).

Although stacking base-level predictions from HMMs with fixed hyperparameters was relatively ineffective for the majority of traits in the HCP dataset, it was extremely effective for one of the subject traits (including when compared to simple averaging). **Figure 3b** shows the accuracy values for all approaches for the selected traits in HCP, highlighting the relative ineffectiveness of stacking for most subject traits. Across all subject traits, stacking performed insignificantly better than base-level predictions (mean R^2^ _STACKED_: 0.0415 vs. mean R^2^ _BASE_: 0.0374; *p_BH_*=0.166); however, the improvement was driven by the specific subject trait ‘PicVocab_Unadj’, where stacking was extremely effective compared to both base-level predictions and averaging (mean R^2^ _STACKED_: 0.114, compared to R^2^ _BASE_: 0.0757, compared to R^2^ _AVERAGE_: 0.0903). For this trait, stacking was very effective due to the high accuracy of a few base-level predictions compared to the remaining predictions, as indicated by the outliers of the boxplot in **Figure 3b**. Although it is visually clear that stacking outperforms simple averaging for this trait, across all traits stacking performed similarly to averaging (mean R^2^ _STACKED_: 0.0415 vs. mean R^2^ _AVERAGE_: 0.0396; *p_BH_*=0.436)

These results highlight the relative effectiveness of stacking in cases where specific random initialisations of the HMM result in significantly more accurate base-level predictions compared to other predictions. Stacking has the ability to disregard the less accurate predictions, where simply averaging across base-level predictions cannot. For the remaining traits, the extremely short boxplots highlight the similarity of prediction accuracies for base-level predictions when the HMM hyperparameters are fixed, resulting in ineffective stacking.

### 3.2 Stacking predictions generated from HMMs with varying hyperparameters improves prediction accuracy to a larger extent

For UKB, stacked predictions obtained from HMMs with varying hyperparameters were generally more accurate than alternative approaches, as shown by **Figure 3b**. Across the 15 traits and 10 cross-validation repetitions, the stacked predictions exhibited significant improvements over the base-level predictions (mean R^2^ _STACKED_: 0.0320 vs. mean R^2^ _BASE_: 0.0254; *p_BH_*=4.59e-08), but performed comparably to averaging the base-level predictions (mean R^2^ _STACKED_: 0.0320 vs. mean R^2^ _AVERAGE_: 0.316; *p_BH_*=0.499). Notably, the stacked prediction consistently outperformed every individual base-level prediction for each subject trait, highlighting the benefit of stacking over selecting a single configuration of the HMM for predictions. Although the improvement of stacking predictions from HMMs with varying hyperparameters over HMMs with fixed hyperparameters was insignificant (mean R^2^ _VARY_: 0.0320 vs. mean R^2^ _FIXED_: 0.0299; *p_BH_*=0.184), the boxplots in **Figure 3b** show a consistent albeit small improvement across subject traits. The left panel of **Figure 3c** shows the difference in these R^2^ scores, highlighting this small but consistent improvement.

For HCP, varying the hyperparameters was also effective. Across the 15 traits and 10 cross-validation repetitions, we found that the stacked predictions outperformed the base-level predictions significantly (mean R^2^ _STACKED_: 0.0438 vs. mean R^2^ _BASE_: 0.0360; *p_BH_*=0.0104) but performed comparably to averaging the base-level predictions (mean R^2^ _STACKED_: 0.0438 vs. mean R^2^ _AVERAGE_: 0.0446; *p_BH_*=0.633). Moreover, stacking predictions from HMMs with varying hyperparameters was only insignificantly better than stacking predictions from HMMs with fixed hyperparameters, as shown by **Figure 3b** and the right panel of **Figure 3c** (mean R^2^ _VARY_: 0.0438 vs. mean R^2^ _FIXED_: 0.0415; *p_BH_*=0.414).

While stacking and simple averaging produced similar results in this specific scenario, it is important to note that this may not always be the case. As illustrated in **Supplementary Figure SI-2**, we replicated the analysis by parametrising the HMM to primarily focus on FC. In this particular configuration, stacking consistently achieved significantly higher results compared to simple averaging in UKB, and yielded much higher accuracy for certain traits in HCP. The circumstances under which stacking outperforms averaging are difficult to anticipate. Therefore, it is generally recommended to employ stacking over averaging, as it is expected to yield at least comparable if not superior performance with little extra computational expense.

### 3.3 Stacking leads to a more robust prediction

While diversity among base-level predictions can result in improved stacked predictions (e.g., through varying the HMM hyperparameters), it is important to ensure that diversity is not driven by a lack of robustness, where model accuracies vary depending on the subjects used for training. To assess the robustness of the predictions when subjects were randomised across cross-validation folds, we conducted 10 cross-validation iterations independently for 50 HMMs with fixed hyperparameters and 50 HMMs with varying hyperparameters.

For UKB, **Figure 4a** compares the variance in accuracy values for the base-level predictions and the stacked predictions across cross-validation iterations (i.e., full repeats of the nested cross-validation framework with different randomised fold distribution). Stacking predictions obtained from HMMs with fixed hyperparameters (light and dark blue boxplots) was ineffective at increasing robustness for any of the subject traits. In contrast, when stacking predictions from HMMs with varying hyperparameters (yellow and gold boxplots), our framework significantly improved robustness across cross-validation iterations for 11 of the 15 traits (p_BH_<0.01^5^).

**Figure 4.**
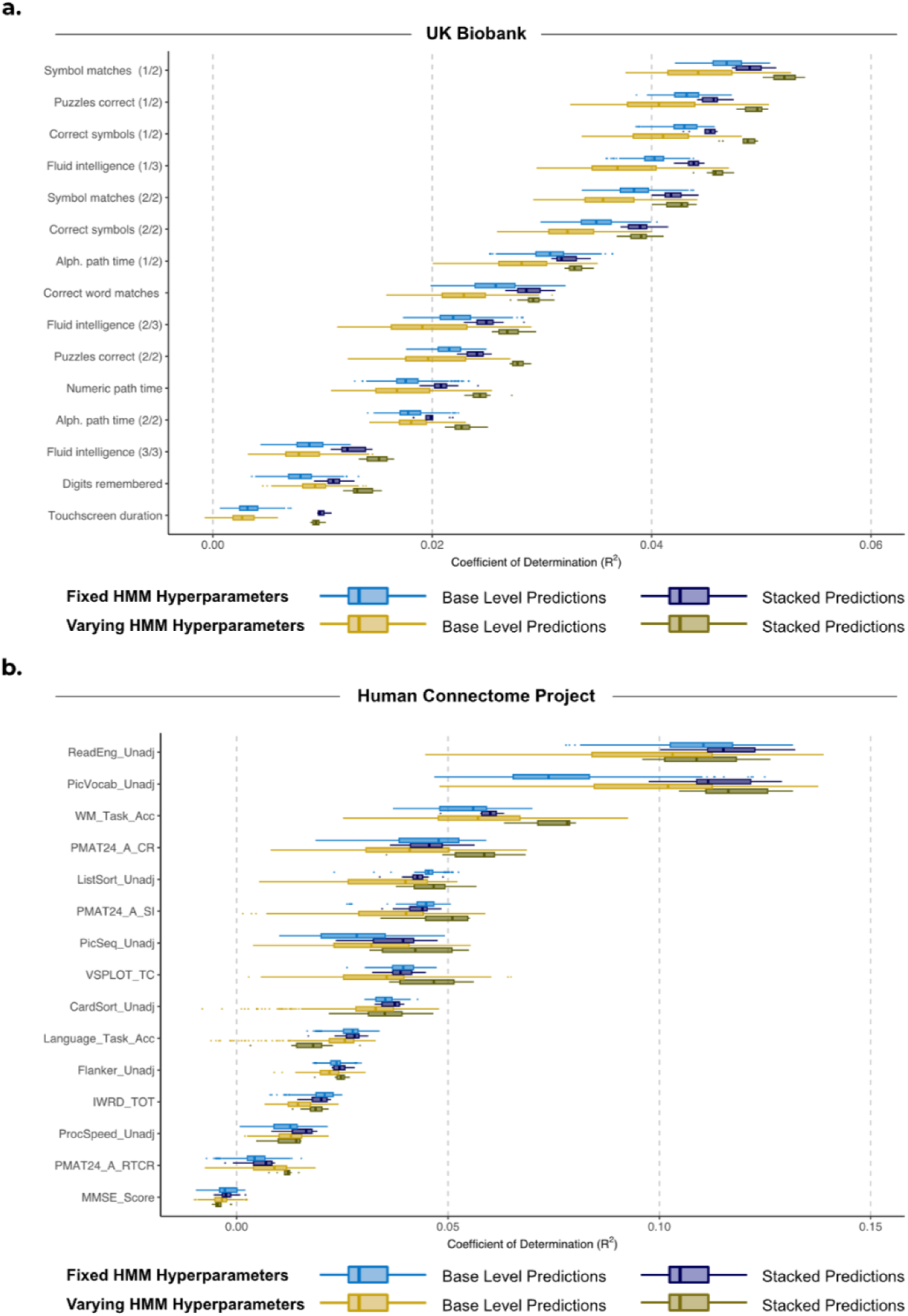
Comparison of variance across 10 cross-validation iterations in performance for base-level predictions and using stacking of HMMs with varying hyperparameters against HMMs with fixed hyperparameters. **(a)** UK Biobank. **(b)** Human Connectome Project. ‘Base-level Predictions’ boxplots (light blue and light yellow) show the R^2^ scores between observed subject traits and 500 base-level predictions (50 HMMs across 10 cross-validation iterations). ‘Stacked Predictions’ boxplots (dark blue and gold) show the R^2^ scores between observed subject traits and 10 stacked predictions (for each cross-validation iteration). Blue represents the results of using HMMs with fixed model hyperparameters. Yellow represents the results of using HMMs with varying model hyperparameters. Stacking was more effective at producing a robust prediction in UKB than HCP and when combining predictions from HMMs with varying hyperparameters.

**Figure 5.**
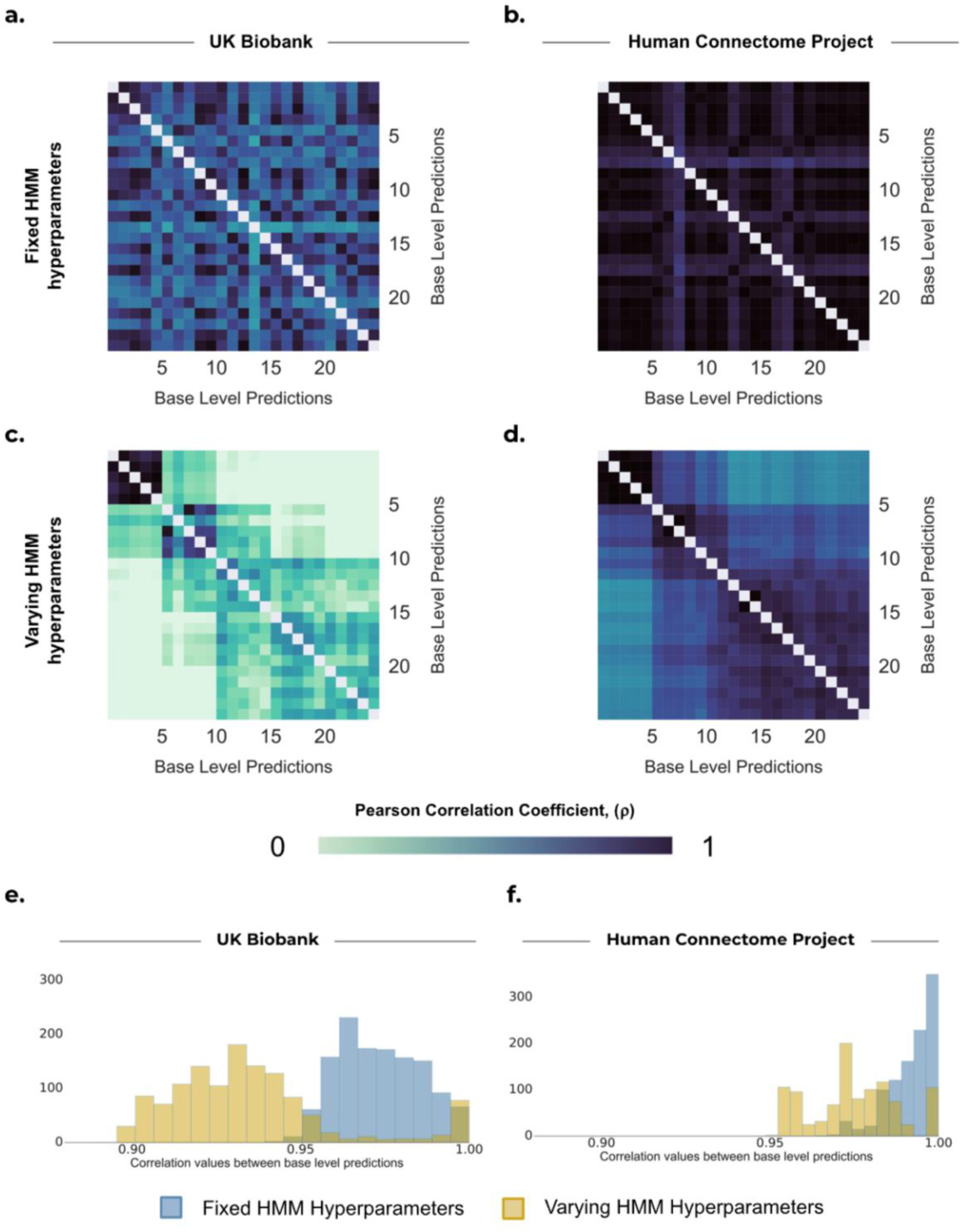
Correlation between base-level predictions for predicting fluid intelligence in UKB and HCP subjects. The correlations between the first 25 of the 50 base-level predictions is shown (i.e., a single run per HMM hyperparameter configuration—see **Supplementary Table SI-3** for more details). **(a) – (d)** Heatmaps show the mean correlation between base-level predictions across 10 cross-validation repetitions obtained from 25 independent runs of the HMM using **(a)** fixed hyperparameters (UKB); **(b)** fixed hyperparameters (HCP); **(c)** varying hyperparameters (UKB); **(d)** varying hyperparameters (HCP). **(e) – (f)** Distribution of off-diagonal values in the correlation matrices. The correlations between base-level predictions are comparatively lower in **(e)** UKB than **(f)** HCP, and varying the hyperparameters leads to even lower correlations. The wider range of accuracies in **(f)** reflects the fact that stacking in HCP is more effective for some traits than others.

**Figure 6.**
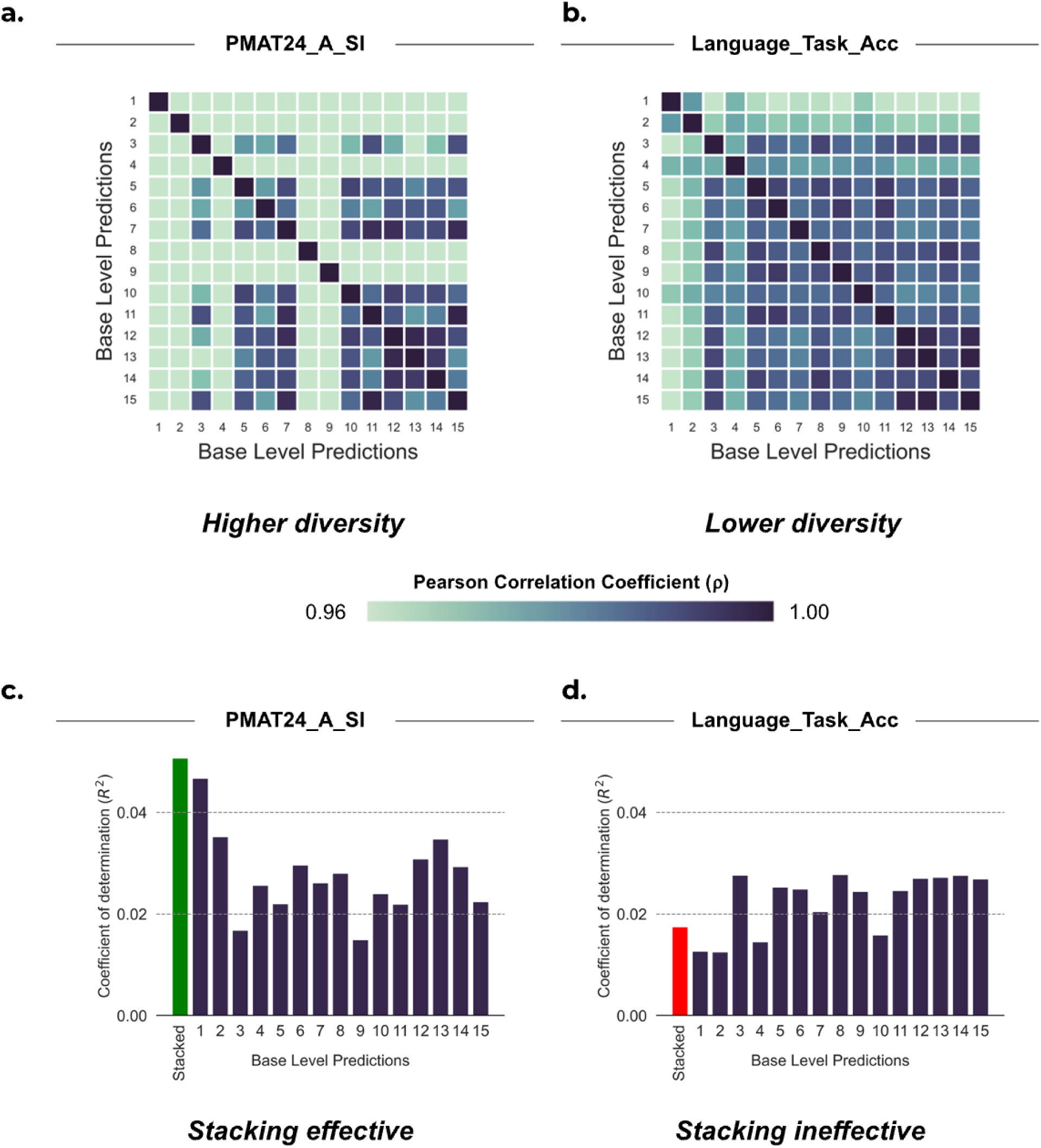
Diversity and accuracy analysis of 15 base-level predictions corresponding to the largest stacking weights from the pool of 50 models for predicting two cognitive traits from HCP. In the top panel, correlation between base-level predictions is shown for **(a)** PMAT24_A_SI and **(b)** Language_Task_Acc. In the bottom panel, the accuracy of base-level predictions (R^2^) is shown for **(c)** PMAT24_A_SI and **(d)** Language_Task_Acc. The predictions were all generated from HMMs with varying hyperparameters. Higher diversity in the base level predictions resulted in more effective stacking for PMAT24_A_SI.

For HCP, stacking was only moderately effective at producing robust predictions, as shown in **Figure 4b**. Stacking predictions obtained from HMMs with fixed hyperparameters (light and dark blue boxplots) significantly increased robustness for only 1 out of 15 subject traits (*p_BH_*<0.01^5^). Moreover, stacking predictions obtained from HMMs with varying hyperparameters (yellow and gold boxplots) increased robustness for 3` out of 15 subject traits (*p_BH_*<0.01^5^).

Overall, stacking predictions demonstrated greater robustness in UKB compared to HCP. This disparity can be attributed to the larger number of subjects in UKB, resulting in stacked predictions which were consistently accurate in UKB. Conversely, the smaller number of subjects in HCP resulted in higher variability across cross-validation iterations.

Furthermore, stacking predictions showed greater robustness when combining predictions from HMMs with varying hyperparameters. This was likely due to the limited run-to-run variability, but increased variance driven by hyperparameter selection variability. In other words, by varying the hyperparameters of the HMM, the accuracy of the base-level predictions varied greatly; however, stacking was able to consistently find an optimal combination of them, particularly in UKB.

### 3.4 Stacking is effective when base-level predictions are diverse yet accurate

But, when is stacking most useful? To effectively stack multiple base-level predictions, the predictions should both be diverse and show a minimum level of accuracy (Hansen & Salamon, 1990). One way to signify diversity is by seeking predictions that are less correlated with each other, indicating differential information (Tumer & Ghosht, 1996). We found that while varying hyperparameters of the HMM generally led to more diversity and subsequently a more accurate stacked prediction, this was not true if the diversity was driven by noise.

We explored the differences in diversity induced by run-to-run and hyperparameter selection variability across datasets. As an illustrative example, **Figure 5** provides a summary of our findings on the impact of both types of variability of the HMM when predicting fluid intelligence for both datasets. We observed that base-level predictions from HMMs with fixed hyperparameters (**Figure 5a** and **Figure 5b**) exhibited higher correlations with each other compared to predictions from HMMs with varying hyperparameters (**Figure 5c** and **Figure 5d**) in both datasets, and that HCP exhibited higher correlations than UKB (UKB: mean *ρ*_FIXED_: 0.973, compared to mean *ρ*_VARYING_: 0.934; HCP: mean *ρ*_FIXED_: 0.993, compared to mean *ρ*_VARYING_: 0.975).

Greater diversity (indicted by lower correlation values) generally led to improved stacked prediction. Also, stacking was more effective in UKB than in HCP, particularly when varying HMM hyperparameters, as discussed in **Sections 0** and **3.2**. Nonetheless, it is important to ensure that this diversity stems from complementary and meaningful patterns in the data rather than inaccurate predictions (i.e., pure noise).

To illustrate this distinction, **Figure 6** considers predictions obtained from HMMs with varying hyperparameters for two cognitive traits in HCP: ‘PMAT24_A_SI’^6^ and ‘Language_Task_Acc’^7^. **Figure 6a** and **Figure 6b** show that the base-level predictions for the PMAT24_A_SI exhibited a lower correlation between each other (mean *ρ*_VSPLOT_OFF_: 0.896) compared to Language_Task_Acc (mean *ρ*_Language_Task_Acc_: 0.981). This greater diversity in base-level predictions resulted in a more effective stacked prediction for PMAT24_A_SI (mean R^2^ _STACK_: 0.0488; mean R^2^ _AVERAGE_: 0.0449; mean R^2^ _BASE_: 0.0362) than for Language_Task_Acc (mean R^2^ _STACK_: 0.0175; mean R^2^ _AVERAGE_: 0.0260; mean R^2^ _BASE_: 0.0238), as shown in **Figure 6c** and **Figure 6d**.

In the left panel, it is evident that the correlations among the base-level predictions for PMAT24_A_SI vary considerably (**Figure 6a)**, exhibiting a range of accuracies (**Figure 6c)**. Conversely, the right panel shows that the chosen base-level predictions (i.e., the most instrumental in forming the stacked prediction) for Language_Task_Acc generally have a higher correlation between predictions. However, predictions 1, 2, and 4 have a comparatively lower correlation with the other predictions, as shown by the light blue columns (or rows) in the heatmap (**Figure 6b**). However, these three base-level predictions also have the lowest accuracy (R^2^) values (**Figure 6d**). In other words, the diversity in the base-level predictions for Language_Task_Acc stems from inaccurate predictions. This suggests that the variability among the base-level predictions for Language_Task_Acc is likely due to estimation noise rather than capturing meaningful complementary patterns in the data.

Moreover, the inefficacy of stacking for Language_Task_Acc is further hindered by the estimation process of stacking weights, which can also introduce statistical noise. This issue arises due to a lack of robustness in HCP predictions, as described in the previous section. Consequently, the stacking process assigns non-zero stacking weights to less accurate predictions (which it should not), caused by certain models performing well during training on specific subjects, but performing poorly on the test set. This resulted in a stacked prediction for Language_Task_Acc that performs worse than many of the base-level predictions.

Overall, these observations emphasise that stacking is ineffective when the variability among base-level predictions is primarily driven by estimation noise rather than capturing distinct and informative aspects of the data.

### 3.5 Dynamic predictions outperform static predictions with sufficient data

While dynamic models, such as the HMMs above, can potentially capture more information, predictions based on static FC may show an advantage due to their simplicity and reliability. Our approach to generating predictions from static FC was chosen to be comparable to our predictions from dynamic FC, given the current prediction algorithm. Furthermore, our goal for this analysis was to investigate if predictions from static FC can complement, or improve upon, predictions from dynamic FC. Given that the case of varying hyperparameter (exhibiting both run-to-run variability and hyperparameter selection variability) subsumes the case of fixed hyperparameters (run-to-run variability only), we therefore focus solely on predictions from dynamic FC generated from HMMs with varying hyperparameters.

For UKB, **Figure 7a** shows that the predictions from static FC performed comparably to the stacked dynamic predictions, which combined base-level predictions from HMMs with varying hyperparameters (mean R^2^ _STATIC_: 0.0323 vs. mean R^2^ _DYNAMIC (STACKED)_: 0.0320; *p_BH_*=0.889). Combining the static and dynamic base-level predictions using our stacking framework (i.e., the 50 dynamic base-level predictions and one prediction from static FC) resulted in a marginal but insignificant improvement in accuracy to stacking only the predictions from dynamic FC (mean R^2^ _STATIC + DYNAMIC (STACKED)_: 0.0340 vs. mean R^2^ _DYNAMIC (STACKED)_: 0.0320; *p_BH_*=0.184) and to averaging across the 51 predictions (mean R^2^ _STATIC + DYNAMIC (STACKED)_: 0.0340 vs. mean R^2^ _STATIC + DYNAMIC (AVERAGE)_: 0.0316; *p_BH_*=0.140).

**Figure 7.**
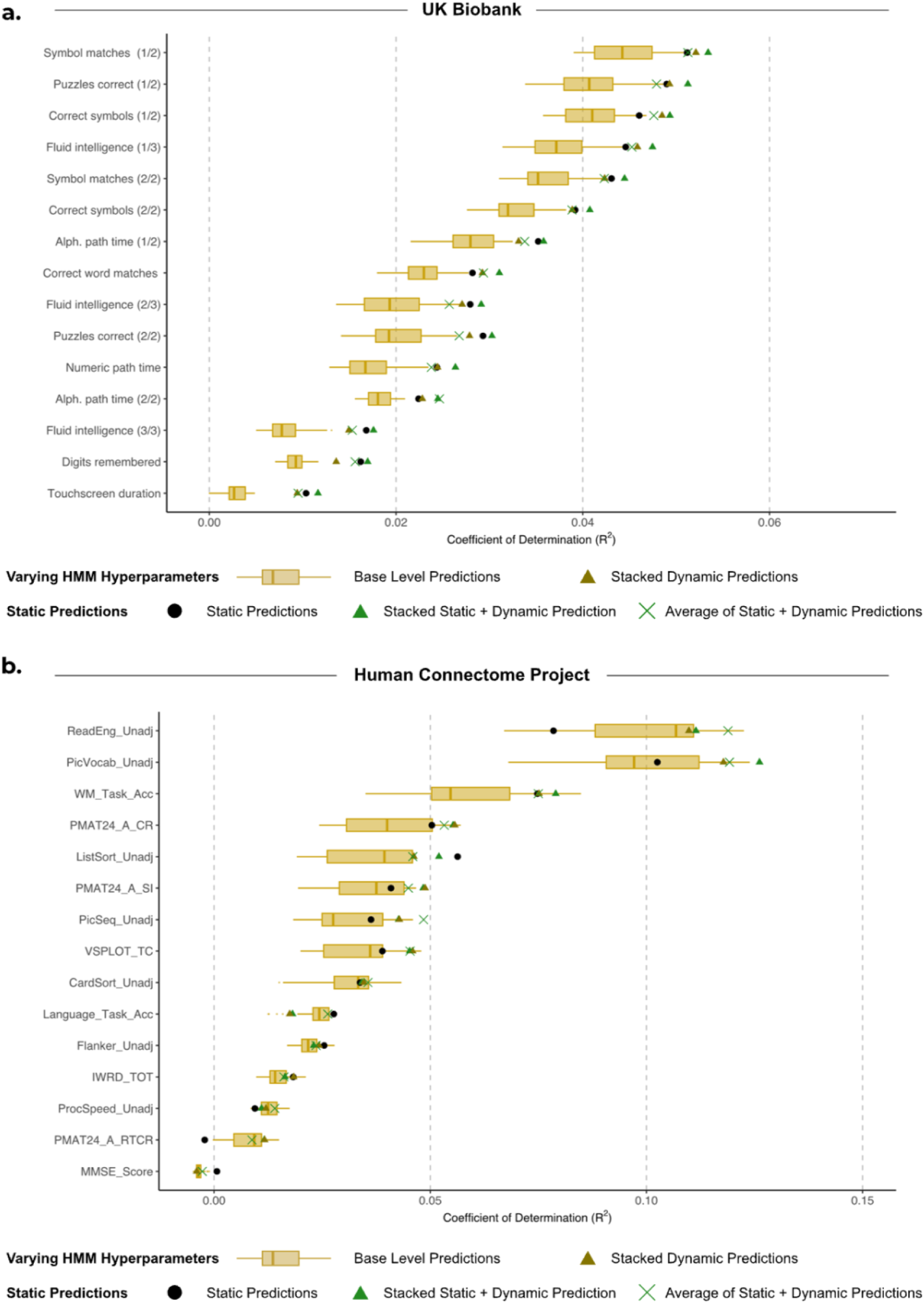
Comparison of performance for static FC, base-level dynamic FC predictions, and stacking the predictions. **(a)** UKB. **(b)** HCP. Boxplots show the R^2^ scores between observed subject traits and dynamic base-level predictions generated from 50 HMMs, and are compared to predictions from static FC (•). These individual predictions are then compared to the R^2^ scores when we combine the base-level predictions by stacking the dynamic base-level predictions with varying HMM hyperparameters (Δ), as well as stacking the predictions from static FC with dynamic base-level predictions (Δ).

For HCP, **Figure 7b** highlights that the predictions from static FC performed poorly for certain subject traits (i.e., the top performing traits; ReadEnd_Unadj and PicVocab_Unadj) compared to predictions from dynamic FC. However, averaged across all subject traits, the improvement of predictions from dynamic FC over static FC was insignificant (mean R^2^ _VARYING (STACKED)_: 0.0438 vs. mean R^2^ _STATIC_: 0.0394; *p_BH_*=0.372). Stacking the dynamic predictions together with the static prediction resulted in comparable accuracies to only stacking the predictions from dynamic FC (mean R^2^ _DYNAMIC (STACKED)_: 0.0447 vs. mean R^2^ _STATIC + DYNAMIC (STACKED)_: 0.0438; *p_BH_*=0.499), and to averaging across dynamic and static predictions (mean R^2^ _DYNAMIC (STACKED)_: 0.0447 vs. mean R^2^ _STATIC + DYNAMIC (AVERAGE)_: 0.0446; *p_BH_*=0.583).

Our findings suggest that dynamic FC tends to generate more accurate predictions than static FC when there is sufficient high-quality data, and for traits that can be predicted to a certain degree of accuracy, as observed in HCP. The HCP dataset, with its 4800 time points compared to UK’s 490, provided ample data for dynamic FC to yield superior prediction accuracy. Conversely, in UKB, where scanning sessions were shorter, the performance of predictions from dynamic FC was comparable to that of static FC for the traits we examined.

As previously discussed, the approach we have focused on for developing predictions from static FC, which used full correlations between brain regions, was specifically chosen to be comparable to our predictions from dynamic FC. In addition, we investigated an alternative approach for developing predictions from static FC, which involved constructing partial correlation network matrices directly from the brain timeseries. The results of this analysis, available in **Supplementary Figure SI-3**, demonstrate the superiority of partial correlations over full correlations.

## 4 Discussion

Generating robust and accurate predictions of subject traits (e.g., clinical or psychological) from brain data is crucial for both the interpretation and the clinical translation of findings. Previous research has shown that dynamic (time-varying) representations of the brain can add information above and beyond that of static (time-averaged) functional representations (Liégeois et al., 2019; Vidaurre et al., 2021), resulting in superior prediction accuracies for behavioural traits using brain dynamics—particularly when combined with the Fisher kernel (Ahrends et al., 2023). To complement structural representations of the brain, and potentially improve over static FC representations, we employed generative models of brain network dynamics and extracted features suitable for subject trait prediction. By combining predictions from multiple brain dynamic models that provided distinct and complementary perspectives of the data, we presented two major benefits of stacking. Firstly, we showed that stacked predictions were more accurate and robust across cross-validation iterations than base-level predictions; this is consistent with previous work on model combination for statistical testing (Vidaurre et al., 2019). Secondly, we highlighted how this approach eliminates the need to arbitrarily select the ‘best’ HMM hyperparameters, and instead incorporates information from many options, ensuring superior predictions across a diverse set of subject traits (albeit at the cost of greater computational expense).

We found that variability in the base-level predictions obtained from HMMs with fixed hyperparameters (i.e., those with run-to-run variability) enhanced the stacked prediction accuracy compared to the base-level predictions for UKB but was generally ineffective for HCP. Additionally, for both datasets, when the HMM hyperparameters were fixed, stacking performed similarly to simple averaging. This outcome is not surprising, as the variation among HMMs with identical hyperparameters but different initialisations is likely to be limited, and driven by the difference in the quality of estimations rather than differences in information content.

We also found that by exploring a range of HMM hyperparameters, we generated diverse base-level predictions that were combined to produce superior stacked predictions. Previous research has shown that the selection of HMM hyperparameters can influence the sensitivity to different temporal scales in FC changes (Ahrends et al., 2022), and this information can be optimally combined for prediction using our approach. We found that base-level predictions obtained from HMMs with varying hyperparameters were found to be less correlated with each other, indicating diversity, and were stacked to produce more accurate predictions. Furthermore, when we repeated the analysis using an alternative parametrisation of the HMM that focused primarily on FC, stacking was particularly effective compared to simple averaging. Overall, these findings indicate that by varying the hyperparameters of the HMM, we go beyond merely reducing noise through combining predictions. Instead, we uncover unique and diverse FC patterns by using different configurations of the dynamic brain model.

It is important to appreciate the differences between the two considered datasets. In UKB, there is a large number of subjects, but a relatively short scanning time per subject, whereas in HCP we have fewer subjects but they underwent longer scanning sessions. This resulted in two differences. Firstly, our predictions in UKB were robust, whereas the accuracy of predictions in HCP was more susceptible to changes in training examples. This finding is consistent with previous research, which has shown that large sample sizes are necessary for replicable brain-phenotype associations, particularly for intelligence traits (Liu et al., 2023). While this limited the robustness of predictions in HCP, stacking still outperformed alternative approaches in both datasets on average. Secondly, the longer scanning session in HCP resulted in dynamic FC adding valuable information beyond that of static FC, whereas they were comparable in UKB. This suggests that in order to reveal unique information beyond that of static FC when using existing dynamic FC techniques, a dataset with sufficient quantity and quality of data per subject, as found in HCP, is more appropriate. Therefore, we suggest that our stacking framework would be particularly beneficial when applied to higher quality datasets.

It is worth mentioning that there are several measures of static FC. Our approach was chosen to be comparable to our predictions from dynamic FC, given the current prediction algorithm. Our results, that time-varying FC outperformed time-averaged FC for cognitive trait predictions in HCP, supports the conclusions in Vidaurre et al., 2021. However, this is not necessarily optimal, as shown in **Supplementary Figure SI-2**. The use of partial correlation network matrices in our alternative approach resulted in improved predictions from static FC, which generally outperformed our predictions from dynamic FC (which do not use precision but covariance matrices). This is not surprising given previous evidence demonstrating the benefit of using partial correlations over full correlations (Pervaiz et al., 2020). Interestingly, the figure also shows the remarkable effectiveness of stacking static FC predictions from partial correlations together with dynamic FC predictions from full correlations. This further highlights the benefits of using stacking when incorporating diverse prediction approaches. A comprehensive investigation of partial and full correlations in the context of static and dynamic FC, as well as their combination through stacking, remains an area for future research. Importantly, in both our approaches, the relationship between static FC (and consequently the dynamic FC, as state covariances include static FC) and non-imaging behavioural measures can be influenced by the cross-subject variations in the spatial arrangement of functional brain regions (Bijsterbosch et al., 2018, 2019).

We have shown stacking to be particularly effective in large-scale datasets such as UKB, where exploring a wide HMM hyperparameter space can be computationally expensive. To ameliorate computation costs, a preliminary exploration of the hyperparameter space could help to identify the key hyperparameters that contribute to an optimal stacked prediction. Given that the HMM has been found to be relatively stable—for example in discovering consistent resting-state topographies across small changes in the number of states (Baker et al., 2014)—using larger differences between HMM hyperparameters could reduce the number of base-level predictions that need to be generated. This drastically reduces the computation time since the fitting of the individual models is the most expensive aspect of the stacking process.

Our proposed method offers a consistent and flexible approach to integrating multiple models of brain dynamics. To further optimise its performance, we suggest diversifying the range of base-level predictions even more. For example, by applying the HMM to different timeseries from the same group of subjects, analyses from multiple brain imaging modalities (e.g., M/EEG) and/or different brain parcellations could be combined. An alternative stacking method, such as random forests, could be used to capture nonlinear relationships between the base-level predictions and subject traits, particularly if different HMM hyperparameters are optimal for different subjects. Additionally, our approach can be readily adapted to different kernel-based prediction methods or to address classification problems, such as classifying patients with brain disorders, by substituting ridge regression with kernel-based classifiers like kernel logistic regression or SVMs.

Overall, this work demonstrates that combining complementary models of brain dynamics allows us to achieve more accurate and robust predictions. The stacking framework we introduced not only provides flexibility for future exploration, but also reduces the risk of poor predictions due to trivial factors such as poor initialisations, incorrect hyperparameter choice, or cross-validation structure.

## Implementation

The code used in this study was implemented in MATLAB 2019a. The MATLAB code for the HMM analysis is publicly available in the repository of the HMM-MAR toolbox at https://github.com/OHBA-analysis/HMM-MAR, and a Python equivalent is publicly available at https://github.com/vidaurre/glhmm/tree/main. The code for constructing the Fisher kernel can be found at https://github.com/OHBA-analysis/HMM-MAR/blob/master/utils/prediction/hmm_kernel.m.

## Acknowledgements

*D. Vidaurre is supported by a Novo Nordisk Foundation Emerging Investigator Fellowship (NNF19OC-0054895), an ERC Starting Grant (ERC-StG-2019-850404), and a DFF Project 1 from the Independent Research Fund of Denmark (2034-00054B)*

*This research has been conducted in part using the UK Biobank Resource under Application Number 8107. We are grateful to UK Biobank for making the data available, and to all UK Biobank study participants, who generously donated their time to make this resource possible. Analysis was carried out on the clusters at the Oxford Biomedical Research Computing (BMRC) facility. BMRC is a joint development between the Wellcome Centre for Human Genetics and the Big Data Institute, supported by Health Data Research UK and the NIHR Oxford Biomedical Research Centre.*

*Data were provided in part by the Human Connectome Project, WU-Minn Consortium (Principal Investigators: David Van Essen and Kamil Ugurbil; 1U54MH091657) funded by the 16 NIH Institutes and Centers that support the NIH Blueprint for Neuroscience Research; and by the McDonnell Center for Systems Neuroscience at Washington University*.

*This research was funded in part by the Wellcome Trust (215573/Z/19/Z). For the purpose of Open Access, the author has applied a CC BY public copyright licence to any Author Accepted Manuscript version arising from this submission*.

*We additionally thank Chetan Gohil and Rukuang Huang for helpful discussions*.

## Footnotes

1 The initial state probabilities, *π*, are excluded from the prediction analysis since they contain minimal informative information for subject trait predictions at the expense of extra computation.

2 In the HMM-MAR MATLAB toolbox and the glhmm Python toolbox, this is specified by the ‘DirichletDiag’/‘dirichlet_diag’ option.

3 *K* = 6, *δ* = 10

4 *K* ∈ {3, 6, 9, 12, 15}, *δ* ∈ {10,100, 1000, 10000, 100000}. See **Supplementary Table SI-3** for full details.

5 See **Supplementary Table SI-5** for p-values for each trait.iterations and 15 cognitive traits for UKB (left) and HCP (right).

6 ‘PMAT24_A_SI’ refers to the HCP trait ‘Penn Matrix Test: Total Skipped Items’, which counts the number of items not presented because the maximum number of errors allowed was reached

7 ‘Language_Task_Acc’ refers to the HCP trait ‘Language Task overall accuracy’, which is the average accuracy from each condition in the language task

## References

Abrol, A., Fu, Z., Du, Y., & Calhoun, V. D. (2019). Multimodal Data Fusion of Deep Learning and Dynamic Functional Connectivity Features to Predict Alzheimer’s Disease Progression. *Proceedings of the Annual International Conference of the IEEE Engineering in Medicine and Biology Society*, EMBS. 10.1109/EMBC.2019.8856500

Ahrends, C., Stevner, A., Pervaiz, U., Kringelbach, M. L., Vuust, P., Woolrich, M. W., & Vidaurre, D. (2022). Data and model considerations for estimating time-varying functional connectivity in fMRI. NeuroImage, 252. 10.1016/j.neuroimage.2022.119026

Ahrends, C., Woolrich, M., & Vidaurre, D. (2023). Predicting individual traits from models of brain dynamics accurately and reliably using the Fisher kernel. BioRxiv. 10.1101/2023.03.02.530638

Alfaro-Almagro, F., Jenkinson, M., Bangerter, N. K., Andersson, J. L. R., Griffanti, L., Douaud, G., Sotiropoulos, S. N., Jbabdi, S., Hernandez-Fernandez, M., Vallee, E., Vidaurre, D., Webster, M., McCarthy, P., Rorden, C., Daducci, A., Alexander, D. C., Zhang, H., Dragonu, I., Matthews, P. M., … Smith, S. M. (2018). Image processing and Quality Control for the first 10,000 brain imaging datasets from UK Biobank. NeuroImage, 166. 10.1016/j.neuroimage.2017.10.034

Alfaro-Almagro, F., McCarthy, P., Afyouni, S., Andersson, J. L. R., Bastiani, M., Miller, K. L., Nichols, T. E., & Smith, S. M. (2021). Confound modelling in UK Biobank brain imaging. NeuroImage, 224. 10.1016/j.neuroimage.2020.117002

Allen, E. A., Damaraju, E., Plis, S. M., Erhardt, E. B., Eichele, T., & Calhoun, V. D. (2014). Tracking whole-brain connectivity dynamics in the resting state. Cerebral Cortex, 24(3). 10.1093/cercor/bhs352

Alonso, S., & Vidaurre, D. (2023). Towards stability of dynamic FC estimates in neuroimaging and electrophysiology: solutions and limits. BioRxiv. 10.1101/2023.01.18.524539

Amari, S., & Nagaoka, H. (2000). Methods of information geometry, volume 191 of Translations of Mathematical Monographs. In American Mathematical Society.

Andersson, J. L. R., Graham, M. S., Drobnjak, I., Zhang, H., Filippini, N., & Bastiani, M. (2017). Towards a comprehensive framework for movement and distortion correction of diffusion MR images: Within volume movement. NeuroImage, 152. 10.1016/j.neuroimage.2017.02.085

Andersson, J. L. R., Graham, M. S., Zsoldos, E., & Sotiropoulos, S. N. (2016). Incorporating outlier detection and replacement into a non-parametric framework for movement and distortion correction of diffusion MR images. NeuroImage, 141. 10.1016/j.neuroimage.2016.06.058

Baum, L. E., Petrie, T., Soules, G., & Weiss, N. (1970). A Maximization Technique Occurring in the Statistical Analysis of Probabilistic Functions of Markov Chains. The Annals of Mathematical Statistics, 41(1). 10.1214/aoms/1177697196

Beckmann, C. F., & Smith, S. M. (2004). Probabilistic Independent Component Analysis for Functional Magnetic Resonance Imaging. IEEE Transactions on Medical Imaging, 23(2). 10.1109/TMI.2003.822821

Benjamini, Y., & Hochberg, Y. (1995). Controlling the False Discovery Rate: A Practical and Powerful Approach to Multiple Testing. Journal of the Royal Statistical Society: Series B (Methodological), 57(1). 10.1111/j.2517-6161.1995.tb02031.x

Bijsterbosch, J. D., Beckmann, C. F., Woolrich, M. W., Smith, S. M., & Harrison, S. J. (2019). The relationship between spatial configuration and functional connectivity of brain regions revisited. ELife, 8. 10.7554/eLife.44890

Bijsterbosch, J. D., Woolrich, M. W., Glasser, M. F., Robinson, E. C., Beckmann, C. F., Van Essen, D. C., Harrison, S. J., & Smith, S. M. (2018). The relationship between spatial configuration and functional connectivity of brain regions. ELife, 7. 10.7554/eLife.32992

Breiman, L. (1996). Stacked regressions. Machine Learning, 24(1), 49–64.

Dinsdale, N. K., Bluemke, E., Sundaresan, V., Jenkinson, M., Smith, S. M., & Namburete, A. I. L. (2022). Challenges for machine learning in clinical translation of big data imaging studies. Neuron, 110(23), 3866–3881. 10.1016/j.neuron.2022.09.012

Džeroski, S., & Ženko, B. (2004). Is combining classifiers with stacking better than selecting the best one? Machine Learning, 54(3). 10.1023/B:MACH.0000015881.36452.6e

Engemann, D. A., Kozynets, O., Sabbagh, D., Lemaitre, G., Varoquaux, G., Liem, F., & Gramfort, A. (2020). Combining magnetoencephalography with magnetic resonance imaging enhances learning of surrogate-biomarkers. ELife, 9. 10.7554/eLife.54055

Griffanti, L., Salimi-Khorshidi, G., Beckmann, C. F., Auerbach, E. J., Douaud, G., Sexton, C. E., Zsoldos, E., Ebmeier, K. P., Filippini, N., Mackay, C. E., Moeller, S., Xu, J., Yacoub, E., Baselli, G., Ugurbil, K., Miller, K. L., & Smith, S. M. (2014). ICA-based artefact removal and accelerated fMRI acquisition for improved resting state network imaging. NeuroImage, 95. 10.1016/j.neuroimage.2014.03.034

Hansen, L. K., & Salamon, P. (1990). Neural Network Ensembles. IEEE Transactions on Pattern Analysis and Machine Intelligence, 12(10). 10.1109/34.58871

He, T., Kong, R., Holmes, A. J., Nguyen, M., Sabuncu, M. R., Eickhoff, S. B., Bzdok, D., Feng, J., & Yeo, B. T. T. (2020). Deep neural networks and kernel regression achieve comparable accuracies for functional connectivity prediction of behavior and demographics. NeuroImage, 206. 10.1016/j.neuroimage.2019.116276

Jaakkola, T., Diekhans, M., & Haussler, D. (2000). A discriminative framework for detecting remote protein homologies. Journal of Computational Biology, 7(1–2), 95–114.

Jaakkola, T., & Haussler, D. (1998). Exploiting generative models in discriminative classifiers. Advances in Neural Information Processing Systems, 11.

Kaufmann, T., van der Meer, D., Doan, N. T., Schwarz, E., Lund, M. J., Agartz, I., Alnæs, D., Barch, D. M., Baur-Streubel, R., Bertolino, A., Bettella, F., Beyer, M. K., Bøen, E., Borgwardt, S., Brandt, C. L., Buitelaar, J., Celius, E. G., Cervenka, S., Conzelmann, A., … Westlye, L. T. (2019). Common brain disorders are associated with heritable patterns of apparent aging of the brain. Nature Neuroscience, 22(10). 10.1038/s41593-019-0471-7

Kavitha, A., Prakash, S. S., Sreeja, P., & Carshia, A. S. (2019). Investigations on the Functional connectivity disruptive patterns of progressive neurodegenerative disorders. *Proceedings of the Annual International Conference of the IEEE Engineering in Medicine and Biology Society*, EMBS. 10.1109/EMBC.2019.8856919

Leonardi, N., & Van De Ville, D. (2015). On spurious and real fluctuations of dynamic functional connectivity during rest. In NeuroImage (Vol. 104). 10.1016/j.neuroimage.2014.09.007

Liégeois, R., Li, J., Kong, R., Orban, C., Van De Ville, D., Ge, T., Sabuncu, M. R., & Yeo, B. T. T. (2019). Resting brain dynamics at different timescales capture distinct aspects of human behavior. Nature Communications, 10(1). 10.1038/s41467-019-10317-7

Liégeois, R., Ziegler, E., Phillips, C., Geurts, P., Gómez, F., Bahri, M. A., Yeo, B. T. T., Soddu, A., Vanhaudenhuyse, A., Laureys, S., & Sepulchre, R. (2016). Cerebral functional connectivity periodically (de)synchronizes with anatomical constraints. Brain Structure and Function, 221(6). 10.1007/s00429-015-1083-y

Lincoln, W. P., & Skrzypekt, J. (1989). Synergy Of Clustering Multiple Back Propagation Networks. Advances in Neural Information Processing Systems 2.

Liu, S., Abdellaoui, A., Verweij, K. J. H., & van Wingen, G. A. (2023). Replicable brain– phenotype associations require large-scale neuroimaging data. Nature Human Behaviour. 10.1038/s41562-023-01642-5

Masaracchia, L., Fredes, F., Woolrich, M. W., & Vidaurre, D. (2023). Dissecting unsupervised learning through hidden Markov modelling in electrophysiological data. BioRxiv. 10.1101/2023.01.19.524547

Mihalik, A., Brudfors, M., Robu, M., Ferreira, F. S., Lin, H., Rau, A., Wu, T., Blumberg, S. B., Kanber, B., Tariq, M., Garcia, M. E., Zor, C., Nikitichev, D. I., Mourão-Miranda, J., & Oxtoby, N. P. (2019). ABCD Neurocognitive Prediction Challenge 2019: Predicting Individual Fluid Intelligence Scores from Structural MRI Using Probabilistic Segmentation and Kernel Ridge Regression. Lecture Notes in Computer Science (Including Subseries Lecture Notes in Artificial Intelligence and Lecture Notes in Bioinformatics), 11791 LNCS. 10.1007/978-3-030-31901-4_16

Mokhtari, F., Akhlaghi, M. I., Simpson, S. L., Wu, G., & Laurienti, P. J. (2019). Sliding window correlation analysis: Modulating window shape for dynamic brain connectivity in resting state. NeuroImage, 189, 655–666. 10.1016/j.neuroimage.2019.02.001

Mourao-Miranda, J., Reinders, A. A. T. S., Rocha-Rego, V., Lappin, J., Rondina, J., Morgan, C., Morgan, K. D., Fearon, P., Jones, P. B., Doody, G. A., Murray, R. M., Kapur, S., & Dazzan, P. (2012). Individualized prediction of illness course at the first psychotic episode: A support vector machine MRI study. Psychological Medicine, 42(5), 1037– 1047. 10.1017/S0033291711002005

Perrone, M. P., & Cooper, L. N. (1992). When Networks Disagree: Ensemble Methods for Technical Report Hybrid Neural Networks Unclassified. Neural Networks for Speech and Image Processing.

Pervaiz, U., Vidaurre, D., Woolrich, M. W., & Smith, S. M. (2020). Optimising network modelling methods for fMRI. NeuroImage, 211. 10.1016/j.neuroimage.2020.116604

Pievani, M., de Haan, W., Wu, T., Seeley, W. W., & Frisoni, G. B. (2011). Functional network disruption in the degenerative dementias. In The Lancet Neurology (Vol. 10, Issue 9). 10.1016/S1474-4422(11)70158-2

Preti, M. G., Bolton, T. A., & Van De Ville, D. (2017). The dynamic functional connectome: State-of-the-art and perspectives. NeuroImage, 160, 41–54. 10.1016/j.neuroimage.2016.12.061

Quinn, A. J., Vidaurre, D., Abeysuriya, R., Becker, R., Nobre, A. C., & Woolrich, M. W. (2018). Task-evoked dynamic network analysis through Hidden Markov Modeling. Frontiers in Neuroscience, 12(AUG). 10.3389/fnins.2018.00603

Sakoğlu, Ü., Pearlson, G. D., Kiehl, K. A., Wang, Y. M., Michael, A. M., & Calhoun, V. D. (2010). A method for evaluating dynamic functional network connectivity and task-modulation: Application to schizophrenia. *Magnetic Resonance Materials in Physics*, Biology and Medicine, 23(5–6). 10.1007/s10334-010-0197-8

Saunders, C., Gammerman, A., & Vovk, V. (1998). Ridge Regression Learning Algorithm in Dual Variables. Proceedings of the 15th International Conference on Machine Learning.

Schölkopf, B., & Smola, A. J. (2001). Learning with Kernels: Support Vector Machines, Regularization, Optimization, and Beyond Adaptive computation and machine learning. In *Learning with Kernels: Support Vector Machines, Regularization*, Optimization, and Beyond Adaptive computation and machine learning.

Sen, B., & Parhi, K. K. (2021). Predicting Biological Gender and Intelligence from fMRI via Dynamic Functional Connectivity. IEEE Transactions on Biomedical Engineering, 68(3). 10.1109/TBME.2020.3011363

Shahab, S., Mulsant, B. H., Levesque, M. L., Calarco, N., Nazeri, A., Wheeler, A. L., Foussias, G., Rajji, T. K., & Voineskos, A. N. (2019). Brain structure, cognition, and brain age in schizophrenia, bipolar disorder, and healthy controls. Neuropsychopharmacology, 44(5). 10.1038/s41386-018-0298-z

Shawe-Taylor, J., Cristianini, N., & others. (2004). Kernel methods for pattern analysis. Cambridge university press.

Smith, S. M., Alfaro-Almagro, F., Miller, K. L., Smith WIN-FMRIB, S., Andersson, J., Clare, S., Douaud, G., Duff, E., Griffanti, L., Hernandez Fernandez, M., Flitney, D., Jbabdi, S., Jenkin-son, M., Johansen-Berg, H., McCarthy, P., Mortimer, D., Salimi-Khorshidi, G., Okell, T., Sotiropoulos, S., … Polimeni, J. (2022). UK Biobank Brain Imaging Documentation UK Biobank Brain Imaging Documentation Contributors to UK Biobank Brain Imaging. http://www.ukbiobank.ac.uk

Smith, S. M., Beckmann, C. F., Andersson, J., Auerbach, E. J., Bijsterbosch, J., Douaud, G., Duff, E., Feinberg, D. A., Griffanti, L., Harms, M. P., Kelly, M., Laumann, T., Miller, K. L., Moeller, S., Petersen, S., Power, J., Salimi-Khorshidi, G., Snyder, A. Z., Vu, A. T., … Glasser, M. F. (2013). Resting-state fMRI in the Human Connectome Project. NeuroImage, 80. 10.1016/j.neuroimage.2013.05.039

Sudlow, C., Gallacher, J., Allen, N., Beral, V., Burton, P., Danesh, J., Downey, P., Elliott, P., Green, J., Landray, M., & others. (2015). UK biobank: an open access resource for identifying the causes of a wide range of complex diseases of middle and old age. PLoS Medicine, 12(3), e1001779.

Tumer, K., & Ghosht, J. (1996). Error Correlation and Error Reduction in Ensemble Classifiers. Connection Science, 8(3–4).

Tumer, K., & Ghosht, J. (1996). Analysis of Decision Boundaries in Linearly Combined Neural Classifiers. Pattern Recognition, 29(2), 341–348.

Vaghari, D., Kabir, E., & Henson, R. N. (2022). Late combination shows that MEG adds to MRI in classifying MCI versus controls. NeuroImage, 252. 10.1016/j.neuroimage.2022.119054

Van Essen, D. C., Smith, S. M., Barch, D. M., Behrens, T. E. J., Yacoub, E., Ugurbil, K., Consortium, W.-M. H. C. P., & others. (2013). The WU-Minn human connectome project: an overview. Neuroimage, 80, 62–79.

Van Essen, D. C., Ugurbil, K., Auerbach, E., Barch, D., Behrens, T. E. J., Bucholz, R., Chang, A., Chen, L., Corbetta, M., Curtiss, S. W., Della Penna, S., Feinberg, D., Glasser, M. F., Harel, N., Heath, A. C., Larson-Prior, L., Marcus, D., Michalareas, G., Moeller, S., … Yacoub, E. (2012). The Human Connectome Project: A data acquisition perspective. In NeuroImage (Vol. 62, Issue 4). 10.1016/j.neuroimage.2012.02.018

Varoquaux, G., Raamana, P. R., Engemann, D. A., Hoyos-Idrobo, A., Schwartz, Y., & Thirion, B. (2017). Assessing and tuning brain decoders: Cross-validation, caveats, and guidelines. NeuroImage, 145. 10.1016/j.neuroimage.2016.10.038

Vidaurre, D., Abeysuriya, R., Becker, R., Quinn, A. J., Alfaro-Almagro, F., Smith, S. M., & Woolrich, M. W. (2018). Discovering dynamic brain networks from big data in rest and task. Neuroimage, 180, 646–656.

Vidaurre, D., Llera, A., Smith, S. M., & Woolrich, M. W. (2021). Behavioural relevance of spontaneous, transient brain network interactions in fMRI. Neuroimage, 229, 117713.

Vidaurre, D., Smith, S. M., & Woolrich, M. W. (2017). Brain network dynamics are hierarchically organized in time. Proceedings of the National Academy of Sciences, 114(48), 12827–12832.

Vidaurre, D., Woolrich, M. W., Winkler, A. M., Karapanagiotidis, T., Smallwood, J., & Nichols, T. E. (2019). Stable between-subject statistical inference from unstable within-subject functional connectivity estimates. Human Brain Mapping, 40(4). 10.1002/hbm.24442

Wolpert, D. H. (1992). Stacked generalization. Neural Networks, 5(2), 241–259.

Woolrich, M. W., Ripley, B. D., Brady, M., & Smith, S. M. (2001). Temporal autocorrelation in univariate linear modeling of FMRI data. NeuroImage, 14(6). 10.1006/nimg.2001.0931

