## Supplementary Material for "Stacking models of brain dynamics improves prediction of subject traits in fMRI"

***Table SI-1*** *HCP Subject Traits.*

| Var. no. | Column Header | Full Display Name | Assessment | HCP Var. no. |
|---|---|---|---|---|
| 1 | ReadEng_Unadj | NIH Toolbox Oral Reading Recognition Test: Unadjusted Scale Score | Language/Reading | 231 |
| 2 | PicVocab_Unadj | NIH Toolbox Picture Vocabulary Test: Unadjusted Scale Score | Language/Vocabulary | 233 |
| 3 | WM_Task_Acc | Working Memory Task OVERALL Accuracy | Working Memory Task | 545 |
| 4 | PMAT24_A_CR | Penn Progressive Matrices: Number of Correct Responses (PMAT24_A_CR) | Fluid Intelligence | 228 |
| 5 | ListSort_Unadj | NIH Toolbox List Sorting Working Memory Test: Unadjusted Scale Score | Working Memory | 264 |
| 6 | PMAT24_A_SI | Penn Progressive Matrices: Total Skipped Items (PMAT24_A_SI) | Fluid Intelligence | 229 |
| 7 | PicSeq_Unadj | NIH Toolbox Picture Sequence Memory Test: Unadjusted Scale Score | Episodic Memory | 222 |
| 8 | VSPLOT_TC | Variable Short Penn Line Orientation: Total Number Correct (VSPLOT_TC) | Spatial Orientation | 251 |
| 9 | CardSort_Unadj | NIH Toolbox Dimensional Change Card Sort Test: Unadjusted Scale Score | Executive Function/ Cognitive Flexibility | 224 |
| 10 | Language_Task_Acc | Language Task OVERALL Accuracy | Language Task | 510 |
| 11 | Flanker_Unadj | NIH Toolbox Flanker Inhibitory Control and Attention Test: Unadjusted Scale Score | Executive Function/ Inhibition | 226 |
| 12 | IRWD_TOT | Penn Word Memory Test: Total Number of Correct Responses (IWRD_TOT) | Verbal Episodic Memory | 262 |
| 13 | ProcSpeed_Unadj | NIH Toolbox Pattern Comparison Processing Speed Test: Unadjusted Scale Score | Processing Speed | 235 |
| 14 | PMAT_24_A_RTCR | Penn Progressive Matrices: Median Reaction Time for Correct Responses (PMAT24_A_RTCR) | Fluid Intelligence | 230 |
| 15 | MMSE_SCORE | Mini Mental Status Exam Total Score | Cognitive Status (Mini Mental Status Exam) | 196 |

***Table SI-2*** *UKB Subject Traits.*

| Var. no. | Column Header | Full Display Name | Category | UKB Field ID |
|---|---|---|---|---|
| 1 | Symbol matches (1/2) | Number of symbol digit matches attempted (2.0) | Symbol digit substitution | 23323 |
| 2 | Puzzles correct (1/2) | Number of puzzles correctly solved (2.0) | Matrix pattern completion | 6373 |
| 3 | Correct symbols (1/2) | Number of symbol digit matches made correctly (2.0) | Symbol digit substitution | 23324 |
| 4 | Fluid intelligence (1/3) | Fluid intelligence score (2.0) | Fluid intelligence / reasoning (cognitive function summary) | 20016 |
| 5 | Symbol matches (2/2) | Number of symbol digit matches made attempted (0.0) | Symbol digit substitution | 23323 |
| 6 | Correct symbols (2/2) | Number of symbol digit matches made correctly (0.0) | Symbol digit substitution | 23324 |
| 7 | Alph. path time (1/2) | Duration to complete alphanumeric path (trail #2) (0.0) | Trail making | 6350 |
| 8 | Correct word matches | Number of word pairs correctly associated (2.0) | Paired associate learning | 20197 |
| 9 | Fluid intelligence (2/3) | Fluid intelligence score (0.0) | Fluid intelligence / reasoning (cognitive function summary) | 20016 |
| 10 | Puzzles correct (2/2) | Number of puzzles correct (2.0) | Tower rearranging | 21004 |
| 11 | Numeric path time | Duration to complete numeric path (trail #1) (2.0) | Trail making | 6348 |
| 12 | Alph. path time (2/2) | Duration to complete alphanumeric path (trail #2) (2.0) | Trail making | 6350 |
| 13 | Fluid intelligence (3/3) | Number of fluid intelligence questions attempted within time limit (2.0) | Fluid intelligence / reasoning (cognitive function summary) | 20128 |
| 14 | Digits remembered | Maximum digits remembered correctly (2.0) | Numeric memory | 4282 |
| 15 | Touchscreen duration | Touchscreen duration (2.0) | Process duration | 21622 |

Notes

(0.0) refers to the initial assessment visit (2006-2010) at which participants were recruited and consent given.

(2.0) refers to the first imaging visit (2014+)

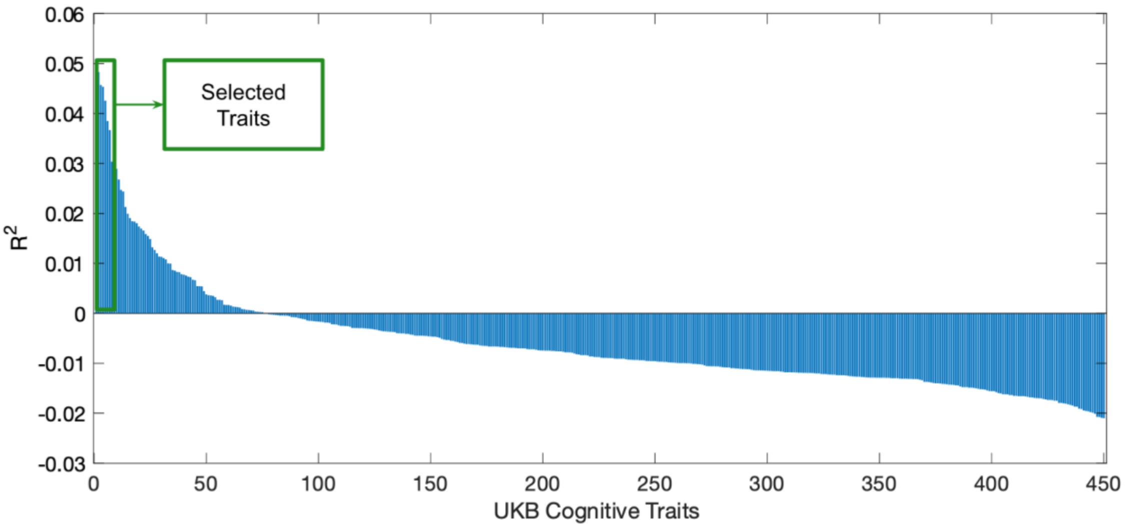


***Figure SI-1*** *Coefficient of determination ($R^2$) between observed subject traits and predictions generated from static FC for the 450 cognitive traits in UKB for which we performed preliminary analysis to select 15 subject traits for further investigation.*

***Table SI-3*** *Choice of HMM hyperparameters for the 50 HMMs where they were varied.*

| HMM run | Number of states ($K$) | Dirichlet distribution concentration parameter ($\delta$) |
|---|---|---|
| 1 | 3 | 10 |
| 2 | 3 | 100 |
| 3 | 3 | 1000 |
| 4 | 3 | 10000 |
| 5 | 3 | 100000 |
| 6 | 6 | 10 |
| 7 | 6 | 100 |
| 8 | 6 | 1000 |
| 9 | 6 | 10000 |
| 10 | 6 | 100000 |
| 11 | 9 | 10 |
| 12 | 9 | 100 |
| 13 | 9 | 1000 |
| 14 | 9 | 10000 |
| 15 | 9 | 100000 |
| 16 | 12 | 10 |
| 17 | 12 | 100 |
| 18 | 12 | 1000 |
| 19 | 12 | 10000 |
| 20 | 12 | 100000 |
| 21 | 15 | 10 |
| 22 | 15 | 100 |
| 23 | 15 | 1000 |
| 24 | 15 | 10000 |
| 25 | 15 | 100000 |
| 26 | 3 | 10 |
| 27 | 3 | 100 |
| 28 | 3 | 1000 |
| 29 | 3 | 10000 |
| 30 | 3 | 100000 |
| 31 | 6 | 10 |
| 32 | 6 | 100 |
| 33 | 6 | 1000 |
| 34 | 6 | 10000 |
| 35 | 6 | 100000 |
| 36 | 9 | 10 |
| 37 | 9 | 100 |
| 38 | 9 | 1000 |
| 39 | 9 | 10000 |
| 40 | 9 | 100000 |
| 41 | 12 | 10 |
| 42 | 12 | 100 |
| 43 | 12 | 1000 |
| 44 | 12 | 10000 |
| 45 | 12 | 100000 |
| 46 | 15 | 10 |
| 47 | 15 | 100 |
| 48 | 15 | 1000 |
| 49 | 15 | 10000 |
| 50 | 15 | 100000 |

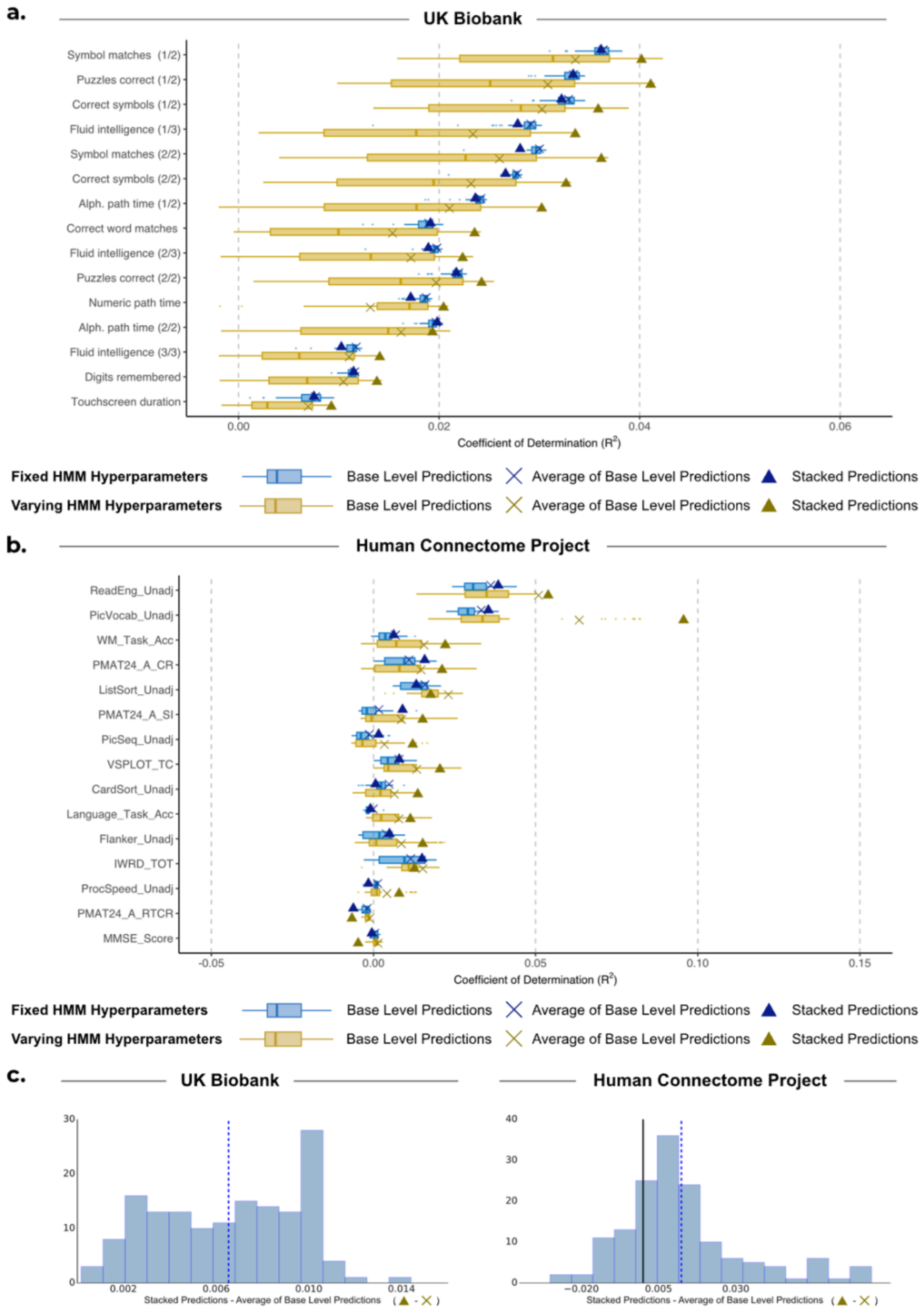


***Figure SI-2** Comparison of performance for base-level predictions and stacking predictions from HMMs with varying hyperparameters against HMMs with fixed hyperparameters, where all HMMs model FC only.* **(a), (b)** Performance of stacking across subject traits for UKB and HCP respectively. Boxplots show the $R^2$ scores between observed subject traits and base-level predictions generated from 50 HMMs. These are compared to the $R^2$ scores when we combine the base-level predictions by taking the average of them (×) and by stacking (▲). Blue represents the results of using HMMs with fixed hyperparameters. Yellow represents the results of using HMMs with varying model hyperparameters. **(c)** Distribution of the difference between stacking predictions (▲) and averaging predictions (×) using varying hyperparameters across 10 cross-validation iterations and 15 cognitive traits for UKB (left) and HCP (right).

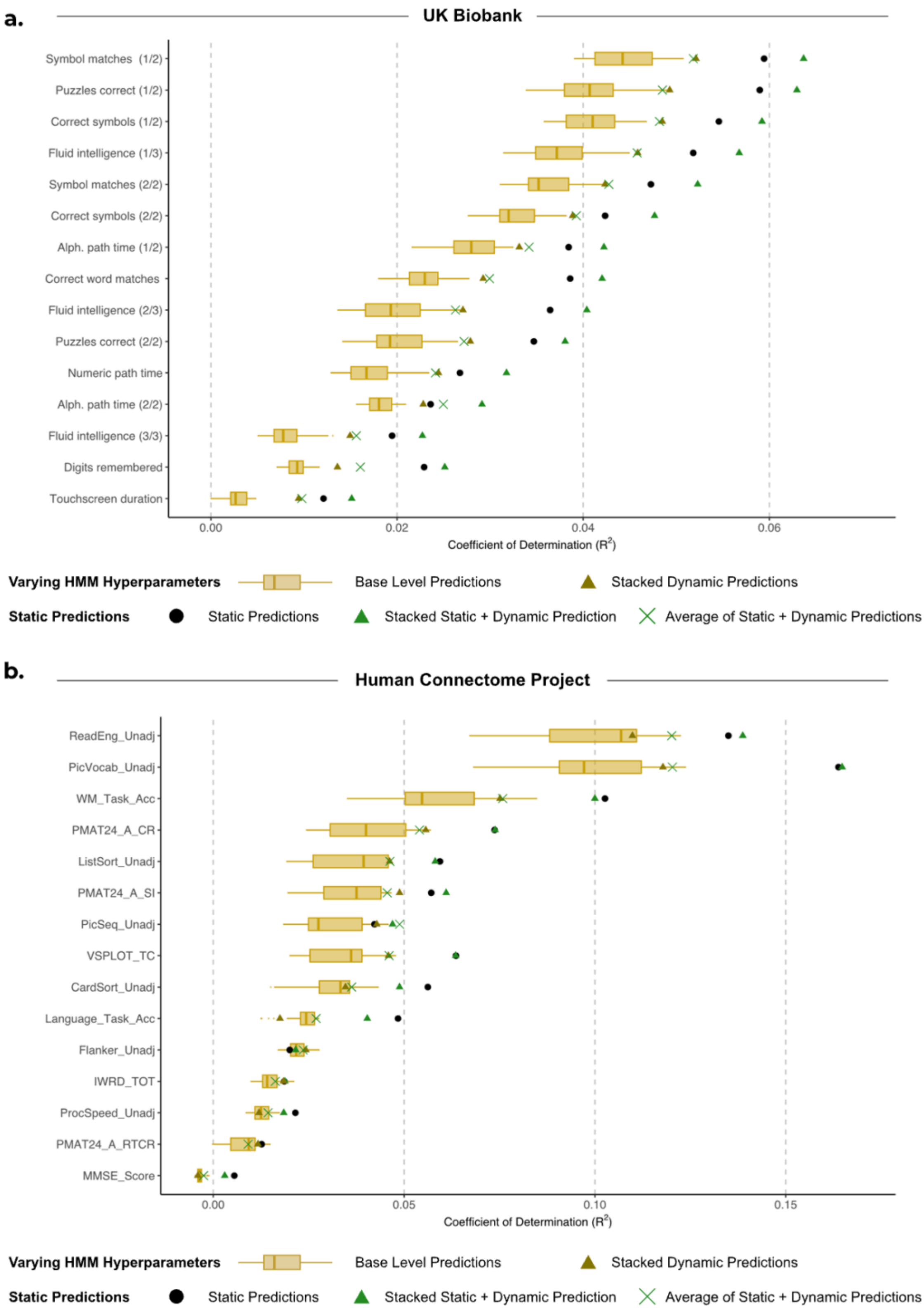


***Figure SI-3*** *Comparison of performance for static FC, base-level dynamic FC predictions, and stacking the predictions. Static predictions have been generated using partial correlation network matrices.* **(a)** UKB. **(b)** HCP. Boxplots show the $R^2$ scores between observed subject traits and dynamic base-level predictions generated from 50 HMMs, and are compared to static FC predictions (●). These individual predictions are then compared to the $R^2$ scores when we combine the base-level predictions by stacking the dynamic base level predictions with varying HMM hyperparameters (▲), as well as stacking the static predictions with dynamic base level predictions (▲). While predictions from static FC using partial correlation network matrices outperform the predictions from dynamic FC using full correlation matrices for both datasets, combining the static and dynamic predictions together results in the most accurate predictions in UKB.

***Table SI-4*** *p-values for Levene's tests comparing the differences in variance between base level predictions generated from HMMs with fixed versus varying hyperparameters for UKB and HCP subject traits. To account for multipe comparisons, we applied Benjamini-Hochberg's False Discovery Rate procedure to correct the p-values within each datset separately. Higher variance across accuracy of base level predictions is an indicator of diversity, which generaly resulted in improved stackde predictions.*

| **Dataset** | **Var. no.** | **Column Header** | **Variance** Fixed HMM Hyperparameters (x e-05) | **Variance** Varying HMM hyperparameters (x e-05) | **p-value** |
|---|---|---|---|---|---|
| UKB | 1 | Symbol matches (1/2) | 0.1325 | 1.333 | 5.85e-09 |
| UKB | 2 | Puzzles correct (1/2) | 0.0960 | 2.006 | 3.08e-08 |
| UKB | 3 | Correct symbols (1/2) | 0.1073 | 0.970 | 8.56e-09 |
| UKB | 4 | Fluid intelligence (1/3) | 0.0927 | 1.730 | 3.08e-08 |
| UKB | 5 | Symbol matches (2/2) | 0.2106 | 0.888 | 1.44e-05 |
| UKB | 6 | Correct symbols (2/2) | 0.2350 | 0.710 | 0.000286 |
| UKB | 7 | Alph. path time (1/2) | 0.2493 | 0.789 | 0.000286 |
| UKB | 8 | Correct word matches | 0.3807 | 0.522 | 0.188 |
| UKB | 9 | Fluid intelligence (2/3) | 0.2230 | 1.654 | 1.76e-08 |
| UKB | 10 | Puzzles correct (2/2) | 0.1181 | 1.206 | 3.08e-08 |
| UKB | 11 | Numeric path time | 0.0987 | 1.034 | 1.36e-07 |
| UKB | 12 | Alph. path time (2/2) | 0.0859 | 0.230 | 0.000312 |
| UKB | 13 | Fluid intelligence (3/3) | 0.1363 | 0.480 | 0.000218 |
| UKB | 14 | Digits remembered | 0.0916 | 0.141 | 0.143 |
| UKB | 15 | Touchscreen duration | 0.0900 | 0.198 | 0.0267 |
| HCP | 1 | ReadEng_Unadj | 0.0371 | 0.0957 | 0.00383 |
| HCP | 2 | PicVocab_Unadj | 0.7354 | 6.6258 | 5.85e-09 |
| HCP | 3 | WM_Task_Acc | 0.0768 | 5.2661 | 1.16e-06 |
| HCP | 4 | PMAT24_A_CR | 0.1196 | 0.901 | 3.08e-06 |
| HCP | 5 | ListSort_Unadj | 0.2902 | 9.7481 | 9.55e-13 |
| HCP | 6 | PMAT24_A_SI | 0.1317 | 7.0784 | 2.03e-09 |
| HCP | 7 | PicSeq_Unadj | 0.2069 | 2.2424 | 4.82e-09 |
| HCP | 8 | VSPLOT_TC | 2.0448 | 31.8699 | 3.08e-08 |
| HCP | 9 | CardSort_Unadj | 8.799 | 28.0428 | 0.000466 |
| HCP | 10 | Language_Task_Acc | 0.1986 | 0.6268 | 0.000524 |
| HCP | 11 | Flanker_Unadj | 0.171 | 6.0937 | 3.65e-11 |
| HCP | 12 | IRWD_TOT | 0.3566 | 0.9865 | 0.0119 |
| HCP | 13 | ProcSpeed_Unadj | 0.0526 | 9.8349 | 3.63e-14 |
| HCP | 14 | PMAT_24_A_RTCR | 0.0613 | 1.2784 | 0.00371 |
| HCP | 15 | MMSE_SCORE | 0.2824 | 11.2517 | 4.87e-07 |

*[ns]not statistically significant results*

***Table SI-5*** *p-values for one-sample Kolmogorov-Smirnov test assessing if the (standardised) distribution of accuracy values ($R^2$; coefficients of determination) for the base-level and stacked predictions for traits in UKB and HCP come from a normal distribution.*

| **Dataset** | **HMM Hyperparameter Type** | **Var. no.** | **Column Header** | **p-vals** Base-Level Prediction | **p-vals** Stacked Prediction |
|---|---|---|---|---|---|
| UKB | Fixed | 1 | Symbol matches (1/2) | 0.4448 | 0.776 |
| UKB | Fixed | 2 | Puzzles correct (1/2) | 0.6784 | 0.8721 |
| UKB | Fixed | 3 | Correct symbols (1/2) | 0.1531 | 0.5489 |
| UKB | Fixed | 4 | Fluid intelligence (1/3) | 0.5895 | 0.9679 |
| UKB | Fixed | 5 | Symbol matches (2/2) | 0.7753 | 0.7004 |
| UKB | Fixed | 6 | Correct symbols (2/2) | 0.7988 | 0.9605 |
| UKB | Fixed | 7 | Alph. path time (1/2) | 0.7191 | 0.6336 |
| UKB | Fixed | 8 | Correct word matches | 0.4915 | 0.9869 |
| UKB | Fixed | 9 | Fluid intelligence (2/3) | 0.0648 | 0.6612 |
| UKB | Fixed | 10 | Puzzles correct (2/2) | 0.838 | 0.9979 |
| UKB | Fixed | 11 | Numeric path time | <0.0001* | 0.8408 |
| UKB | Fixed | 12 | Alph. path time (2/2) | 0.0097 | 0.5217 |
| UKB | Fixed | 13 | Fluid intelligence (3/3) | 0.8622 | 0.8217 |
| UKB | Fixed | 14 | Digits remembered | 0.8163 | 0.9993 |
| UKB | Fixed | 15 | Touchscreen duration | 0.0618 | 0.7377 |
| UKB | Vary | 1 | Symbol matches (1/2) | 0.0554 | 0.95 |
| UKB | Vary | 2 | Puzzles correct (1/2) | 0.0076 | 0.7784 |
| UKB | Vary | 3 | Correct symbols (1/2) | 0.1641 | 0.541 |
| UKB | Vary | 4 | Fluid intelligence (1/3) | 0.0011* | 0.8222 |
| UKB | Vary | 5 | Symbol matches (2/2) | 0.001* | 0.9341 |
| UKB | Vary | 6 | Correct symbols (2/2) | 0.006* | 0.762 |
| UKB | Vary | 7 | Alph. path time (1/2) | 0.0532 | 0.8707 |
| UKB | Vary | 8 | Correct word matches | 0.259 | 0.9826 |
| UKB | Vary | 9 | Fluid intelligence (2/3) | 0.006* | 0.8914 |
| UKB | Vary | 10 | Puzzles correct (2/2) | 0.0011* | 0.9814 |
| UKB | Vary | 11 | Numeric path time | 0.0017* | 0.9627 |
| UKB | Vary | 12 | Alph. path time (2/2) | 0.3609 | 0.982 |
| UKB | Vary | 13 | Fluid intelligence (3/3) | 0.0007* | 0.9579 |
| UKB | Vary | 14 | Digits remembered | 0.5598 | 0.7284 |
| UKB | Vary | 15 | Touchscreen duration | 0.396 | 0.9674 |
| HCP | Fixed | 1 | ReadEng_Unadj | <0.0001* | 0.8496 |
| HCP | Fixed | 2 | PicVocab_Unadj | 0.0042* | 0.8183 |
| HCP | Fixed | 3 | WM_Task_Acc | 0.0062* | 0.7994 |
| HCP | Fixed | 4 | PMAT24_A_CR | 0.0014* | 0.9668 |
| HCP | Fixed | 5 | ListSort_Unadj | <0.0001* | 0.9645 |
| HCP | Fixed | 6 | PMAT24_A_SI | <0.0001* | 0.6162 |
| HCP | Fixed | 7 | PicSeq_Unadj | <0.0001* | 0.5139 |
| HCP | Fixed | 8 | VSPLOT_TC | 0.009* | 0.9471 |
| HCP | Fixed | 9 | CardSort_Unadj | 0.0012* | 0.7162 |
| HCP | Fixed | 10 | Language_Task_Acc | <0.0001* | 0.7765 |
| HCP | Fixed | 11 | Flanker_Unadj | 0.1245 | 0.9184 |
| HCP | Fixed | 12 | IRWD_TOT | 0.0043* | 0.8959 |
| HCP | Fixed | 13 | ProcSpeed_Unadj | <0.0001* | 0.6316 |
| HCP | Fixed | 14 | PMAT_24_A_RTCR | 0.003* | 0.7453 |
| HCP | Fixed | 15 | MMSE_SCORE | 0.005* | 0.3752 |
| HCP | Vary | 1 | ReadEng_Unadj | 0.0212* | 0.6563 |
| HCP | Vary | 2 | PicVocab_Unadj | 0.015* | 0.9213 |
| HCP | Vary | 3 | WM_Task_Acc | <0.0001* | 0.9608 |
| HCP | Vary | 4 | PMAT24_A_CR | 0.0787 | 0.6251 |
| HCP | Vary | 5 | ListSort_Unadj | <0.0001* | 0.8127 |
| HCP | Vary | 6 | PMAT24_A_SI | <0.0001* | 0.5418 |
| HCP | Vary | 7 | PicSeq_Unadj | <0.0001* | 0.6412 |
| HCP | Vary | 8 | VSPLOT_TC | <0.0001* | 0.8011 |
| HCP | Vary | 9 | CardSort_Unadj | <0.0001* | 0.9712 |
| HCP | Vary | 10 | Language_Task_Acc | 0.3198 | 0.2797 |
| HCP | Vary | 11 | Flanker_Unadj | <0.0001* | 0.923 |
| HCP | Vary | 12 | IRWD_TOT | 0.0178* | 0.9667 |
| HCP | Vary | 13 | ProcSpeed_Unadj | <0.0001* | 0.9838 |
| HCP | Vary | 14 | PMAT_24_A_RTCR | <0.0001* | 0.851 |
| HCP | Vary | 15 | MMSE_SCORE | 0.145 | 0.2992 |

**statistically significant results*

***Table SI-6** p-values for Levene's tests comparing the differences in variance between base level predictions and stacked prediction for UKB and HCP subject traits. To account for multipe comparisons, we applied Benjamini-Hochberg's False Discovery Rate procedure to correct the p-values within each datset separately.*

| **Dataset** | **HMM Hyperparameter Type** | **Var. no.** | **Column Header** | **Variance** Base-Level Prediction (x e-05) | **Variance** Stacked Prediction (x e-05) | **p-value** |
|---|---|---|---|---|---|---|
| UKB | Fixed | 1 | Symbol matches (1/2) | 0.1381 | 0.0196 | 0.043* |
| UKB | Fixed | 2 | Puzzles correct (1/2) | 0.2284 | 0.1184 | 0.294 |
| UKB | Fixed | 3 | Correct symbols (1/2) | 0.3199 | 0.2285 | 0.549 |
| UKB | Fixed | 4 | Fluid intelligence (1/3) | 0.2036 | 0.1378 | 0.399 |
| UKB | Fixed | 5 | Symbol matches (2/2) | 0.2206 | 0.1134 | 0.333 |
| UKB | Fixed | 6 | Correct symbols (2/2) | 0.1827 | 0.0662 | 0.231 |
| UKB | Fixed | 7 | Alph. path time (1/2) | 0.2663 | 0.1977 | 0.615 |
| UKB | Fixed | 8 | Correct word matches | 0.4214 | 0.1564 | 0.299 |
| UKB | Fixed | 9 | Fluid intelligence (2/3) | 0.3668 | 0.1681 | 0.231 |
| UKB | Fixed | 10 | Puzzles correct (2/2) | 0.4115 | 0.2370 | 0.275 |
| UKB | Fixed | 11 | Numeric path time | 0.3579 | 0.1772 | 0.346 |
| UKB | Fixed | 12 | Alph. path time (2/2) | 0.6276 | 0.2453 | 0.231 |
| UKB | Fixed | 13 | Fluid intelligence (3/3) | 0.1943 | 0.1210 | 0.459 |
| UKB | Fixed | 14 | Digits remembered | 0.3085 | 0.2032 | 0.615 |
| UKB | Fixed | 15 | Touchscreen duration | 0.2217 | 0.1113 | 0.275 |
| UKB | Vary | 1 | Symbol matches (1/2) | 0.0242 | 0.0020 | 0.030* |
| UKB | Vary | 2 | Puzzles correct (1/2) | 0.0272 | 0.0133 | 0.440 |
| UKB | Vary | 3 | Correct symbols (1/2) | 0.1184 | 0.0161 | 0.013* |
| UKB | Vary | 4 | Fluid intelligence (1/3) | 0.0336 | 0.0121 | 0.160 |
| UKB | Vary | 5 | Symbol matches (2/2) | 0.2086 | 0.0108 | 0.009* |
| UKB | Vary | 6 | Correct symbols (2/2) | 0.1800 | 0.0101 | 0.007* |
| UKB | Vary | 7 | Alph. path time (1/2) | 0.0593 | 0.0128 | 0.089 |
| UKB | Vary | 8 | Correct word matches | 0.0889 | 0.0076 | 0.011* |
| UKB | Vary | 9 | Fluid intelligence (2/3) | 0.0822 | 0.0165 | 0.039* |
| UKB | Vary | 10 | Puzzles correct (2/2) | 0.1793 | 0.0158 | 0.007* |
| UKB | Vary | 11 | Numeric path time | 0.1026 | 0.0208 | 0.059 |
| UKB | Vary | 12 | Alph. path time (2/2) | 0.0707 | 0.0161 | 0.039* |
| UKB | Vary | 13 | Fluid intelligence (3/3) | 0.1253 | 0.0042 | 0.003* |
| UKB | Vary | 14 | Digits remembered | 0.1466 | 0.0188 | 0.012* |
| UKB | Vary | 15 | Touchscreen duration | 0.1077 | 0.0151 | 0.009* |
| HCP | Fixed | 1 | PicVocab_AgeAdj | 0.005 | 0.005 | 0.615 |
| HCP | Fixed | 2 | ReadEng_Unadj | 0.079 | 0.055 | 0.343 |
| HCP | Fixed | 3 | ReadEng_AgeAdj | 0.005 | 0.006 | 0.598 |
| HCP | Fixed | 4 | PicVocab_Unadj | 0.004 | 0.003 | 0.839 |
| HCP | Fixed | 5 | WM_Task_Acc | 0.070 | 0.036 | 0.184 |
| HCP | Fixed | 6 | PMAT24_A_CR | 0.011 | 0.019 | 0.393 |
| HCP | Fixed | 7 | Relational_Task_Acc | 0.016 | 0.017 | 0.945 |
| HCP | Fixed | 8 | ListSort_Unadj | 0.106 | 0.083 | 0.527 |
| HCP | Fixed | 9 | ListSort_AgeAdj | 0.191 | 0.097 | 0.398 |
| HCP | Fixed | 10 | PicSeq_AgeAdj | 0.015 | 0.012 | 0.577 |
| HCP | Fixed | 11 | PicSeq_Unadj | 0.009 | 0.013 | 0.862 |
| HCP | Fixed | 12 | VSPLOT_TC | 0.009 | 0.007 | 0.615 |
| HCP | Fixed | 13 | VSPLOT_OFF | 0.004 | 0.016 | 0.092 |
| HCP | Fixed | 14 | PMAT24_A_SI | 0.011 | 0.017 | 0.667 |
| HCP | Fixed | 15 | Language_Task_Acc | 0.054 | 0.018 | 0.039* |
| HCP | Vary | 1 | PicVocab_AgeAdj | 0.006 | 0.003 | 0.284 |
| HCP | Vary | 2 | ReadEng_Unadj | 0.118 | 0.078 | 0.422 |
| HCP | Vary | 3 | ReadEng_AgeAdj | 0.076 | 0.05 | 0.615 |
| HCP | Vary | 4 | PicVocab_Unadj | 0.012 | 0.005 | 0.177 |
| HCP | Vary | 5 | WM_Task_Acc | 0.162 | 0.084 | 0.171 |
| HCP | Vary | 6 | PMAT24_A_CR | 0.111 | 0.054 | 0.266 |
| HCP | Vary | 7 | Relational_Task_Acc | 0.030 | 0.004 | 0.012* |
| HCP | Vary | 8 | ListSort_Unadj | 0.403 | 0.103 | 0.108 |
| HCP | Vary | 9 | ListSort_AgeAdj | 0.357 | 0.089 | 0.069 |
| HCP | Vary | 10 | PicSeq_AgeAdj | 0.015 | 0.016 | 0.901 |
| HCP | Vary | 11 | PicSeq_Unadj | 0.103 | 0.056 | 0.500 |
| HCP | Vary | 12 | VSPLOT_TC | 0.014 | 0.007 | 0.284 |
| HCP | Vary | 13 | VSPLOT_OFF | 0.125 | 0.037 | 0.033* |
| HCP | Vary | 14 | PMAT24_A_SI | 0.042 | 0.047 | 0.933 |
| HCP | Vary | 15 | Language_Task_Acc | 0.148 | 0.033 | 0.049* |

**statistically significant results*
